# Latent Dirichlet Allocation Mixture Models for Nucleotide Sequence Analysis

**DOI:** 10.1101/2023.12.10.571018

**Authors:** Bixuan Wang, Stephen M. Mount

## Abstract

Strings of nucleotides that carry biological information are typically described using sequence motifs that can be represented by weight matrices or consensus sequences. However, many biological signals in DNA or RNA are recognized by multiple factors in temporal sequence, consist of a mixture of sometimes dissimilar alternative motifs, or may be described better by base composition. Here we apply the Latent Dirichlet Allocation (LDA) mixture model to nucleotide sequences, using k-mers as features, in three related approaches. First, positions in aligned sequences are used as samples. Alternatively, whole sequences are used as samples, either with positional k-mers or with bulk k-mers occurring throughout the sequence. LDA readily identifies motifs, including such elusive cases as the intron branch site. LDA can also identify subtypes of sequence, such as splice site subtypes enriched in long vs. short introns, and can reliably distinguish such properties as reading frame or species of origin. Our results show that LDA is a useful model for describing heterogeneous signals, for assigning individual sequences to subtypes, and for identifying and characterizing sequences that do not fit recognized subtypes. Because LDA topic models are interpretable, they also aid the discovery of new motifs, even those present in a small fraction of samples, allowing a user to hypothesize potential regulatory factors. In summary, LDA can identify and characterize signals in nucleotide sequences and is useful for identifying candidate regulatory factors involved in complex biological processes.

## Introduction

Nucleotide sequences carry information that encodes the complexity and diversity of life. Information in these sequences directs gene expression, from chromatin structure, transcription, splicing, and other steps of RNA processing, to mRNA translation, stability, and localization. In general, signals for these processes are strings of nucleotides that are recognized by RNA or protein molecules, either alone or as part of a multi-subunit complex. These signals are often represented by consensus sequences, nucleotide frequency matrices, position weight matrices (Stormo *et al*. 2000), or Hidden Markov models (Yoon *et al*. 2009), and there are many algorithms for identifying such signals (*e.g.* MEME and its many extensions, Bailey *et al*. 2015). The paradigm is that a signal can be represented by a sequence motif, which functions as the binding site for a single protein or RNA. However, many signals do not fit this paradigm. Some, such as core splice sites) are recognized by multiple factors in a temporal sequence. Others (such as exonic splicing enhancers) may consist of a mixture of alternative motifs recognized by alternative factors, which can differ significantly, or may be completely unrelated. There are also signals that are best described by sequence composition, or by the presence, or absence, of very short motifs. Conversely, even when a single factor is necessary for a process and has a well-defined recognition motif, there are sometimes specific sequences that function without that motif. One example is the general transcription factor TBP, which is thought to function at all transcription events and recognizes a well-defined TATA motif, but not all promoters have this motif (Akhtar & Veenstra, 2011; Cooper et al., 2006). Another example is the branch site, which is recognized by U2 snRNA during the splicing process and has a well-defined sequence motif complementary to U2 (Konarska et al., 1985). Here too, individual introns often lack this motif.

One sequence element that comes in distinct subtypes is the 3’ splice site (Wilkinson et al., 2020). There are three components of the core 3’ splice site: the site itself, including an AG dinucleotide; a pyrimidine tract upstream of position -4; and the branch site, where lariat formation occurs during splicing. In addition, auxiliary sequences such as exonic splicing enhancers in the exon downstream of the splice site contribute to the recognition of 3’ splice sites. The diversity of 3’ splice sites is supported by a variety of observations. Individual human introns differ regarding whether they contain a recognizable branch site motif. While many contain a perfect match to the CTR<u>A</u>Y consensus, others do not, and even the A residue is not invariant at actual sites of branching (Taggart et al., 2017; Wallace & Edmonds, 1983). In addition, 3’ splice sites in short Drosophila introns are functionally distinct from 3’ splice sites in long Drosophila introns (Guo & Mount, 1995).

Latent Dirichlet Allocation (LDA, (Blei et al., 2003)) uses the observation of words in documents to describe the composition of documents in terms of topics that use those words. LDA allows a single sample to have partial membership in each topic, making it possible to recognize similarities between parts of samples (Fig 1A). The topics are matrices of words that are observed, and each individual document is described as a mixture of topics whose fractional representation sums to 1. Documents can be compared by analyzing their topics, and the distinguishing features of each document are interpretable because the driving features in topics are different. LDA has proven to be a useful and extremely popular model in population genetics (Pritchard et al., 2000), where genetic similarity among individuals, geographic isolation, and admixture can be recognized using multi-locus genotype data. Moreover, LDA has been applied to RNA-seq data analysis (Dey et al., 2017), where modeling gene expression data from tissue samples allowed similarities between tissues and estimates of the proportional representation of samples to be described in terms of robust interpretable features that could be easily understood as cell type-specific genes. Those applications of LDA focus on genotype- and gene-level data and show that the clustering power and interpretable features of LDA are useful for describing gene expression data. LDA has also been applied to mutational signatures in cancer (Matsutani et al., 2019), but not to nucleotide sequences more generally.

**Figure 1.**
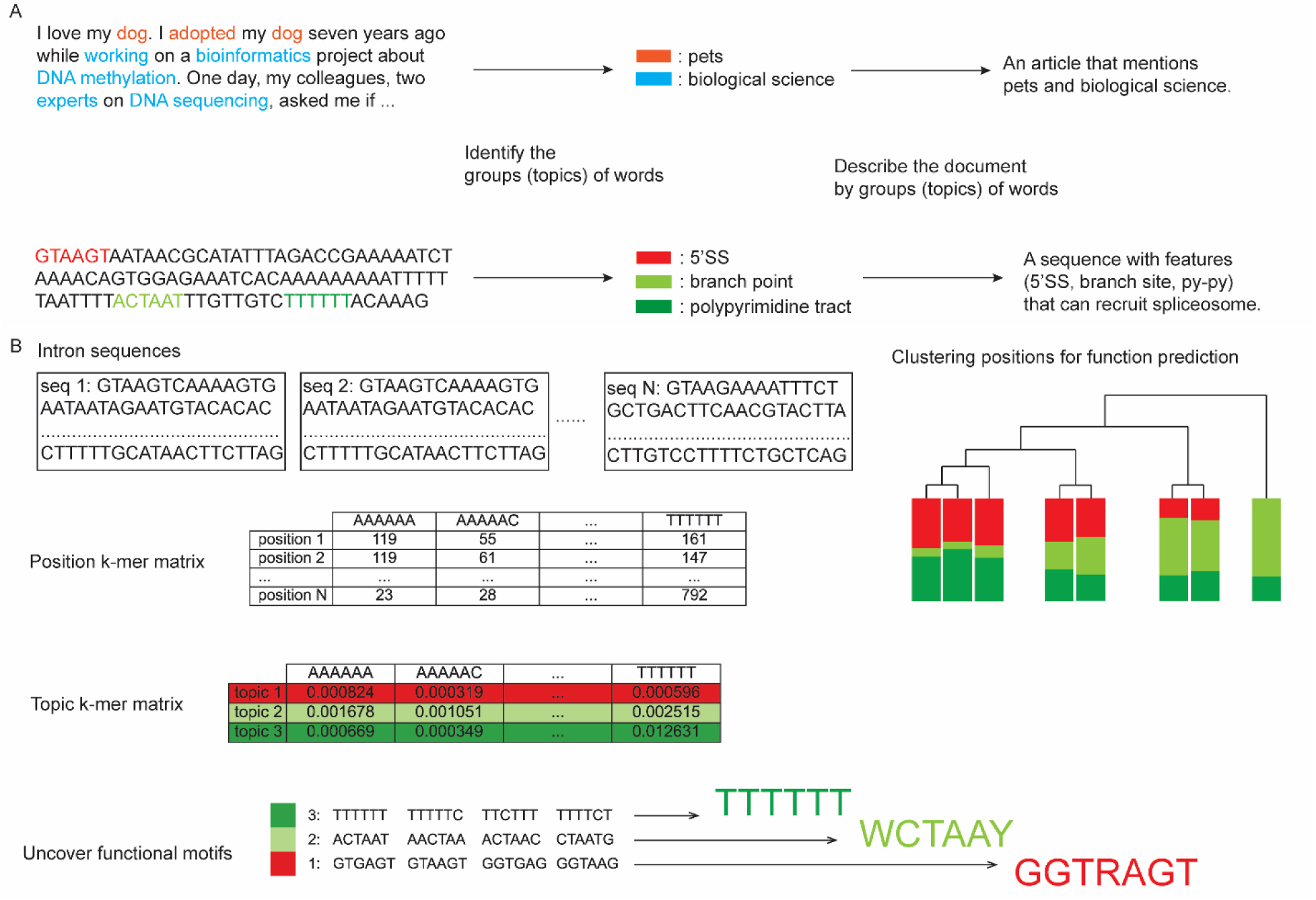
LDA applications in natural language and biological sequences. A. Summarizing topics from input natural language sequences and biological sequences by LDA. LDA can analyze the building blocks from the input sequences (words or nucleotide k-mers) to recognize topics, which describe the features of the input sequences. B. The pipeline of analyzing intron sequence features by LDA. After summarizing the k-mer counts at each position in a matrix, LDA calculates topic k-mer matrices and transforms sequences into topic memberships. Sequence clustering can be achieved by analyzing the topic distributions and the interpretation of topics can reveal functional motifs.

Here we apply the LDA mixture model to nucleotide sequences using k-mers, which are strings of k consecutive nucleotides, as features (Fig 1B). In our first approach, modeling positions in an alignment, sequences are aligned relative to a fixed position such as a splice site, transcription start site or polyadenylation site, and individual positions are used as samples. In a second approach, aligned sequences are used as samples, and positional k-mers, in which the location of the k-mer relative to the point of alignment is part of the feature, are used as features. Finally, a third approach uses k-mer counts in bulk (unaligned) sequences.

Applying LDA to splice sites from introns of different lengths allowed us to clearly identify known core splice site motifs, including the branch site consensus, which is nearly impossible to derive from sequence alone using established methods such as MEME (Bailey et al., 2006). We also identify distinct intron subtypes distinguished by distinct sequence preferences throughout the intron. While these subtypes are preferentially associated with short or long introns there are apparently some long introns of the short subtype and *vice versa*. We also find that LDA can reliably identify the reading frame within coding sequences and can also distinguish human and Drosophila coding sequences. In summary, our results show that LDA is a useful model for describing heterogeneous signals, for assigning individual sequences to subtypes and for identifying sequences that are unusual and do not fit the recognized subtypes.

## Results

### Modeling positions in an alignment identifies signals associated with RNA splicing

To evaluate the ability of mixture models to identify position-related features in batched sequences, we aligned human and Drosophila intron sequences at annotated 3’ splice sites and used k-mer compositions at each position in the alignment (Fig 2A). For the example shown in Fig. 1A, we used 6-mer features, applied a sliding window of six nucleotides, and fitted the LDA model with 6 topics to compare long and short human introns (Fig 2B). The driving k-mers of each topic were obtained based on K-L divergence (as in (Dey et al., 2017)). Examination of the driving k-mers (Fig. 2C) shows that topic 4 contains features similar to the branch site (CTNA), and topic 2 resembles the polypyrimidine tract.

**Figure 2.**
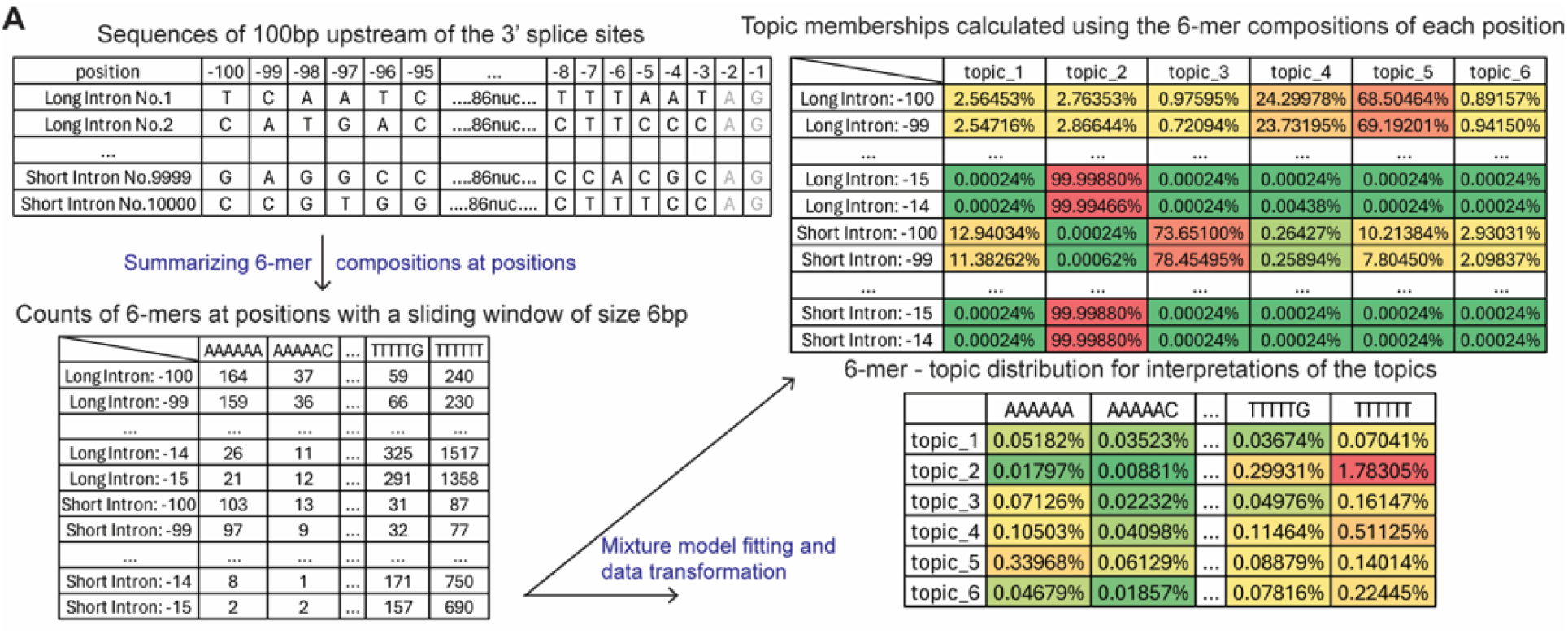
LDA characterization of short and long human introns using aligned positions as samples. LDA was applied to samples corresponding to positions in an alignment of 3’ splice sites from long (>=150 nt.) and short (<150 nt.) introns. A. Diagram of the LDA analysis of intron sequences aligned at the 3’ splice sites. First, sequences are aligned at the 3’ splice sites (up left), and the positions of A and G are -2 and -1. The region from position -100 to -3 are used for the analysis. Then the counts of 6-mers with a sliding window of size 6 at every position are summarized (down left). LDA fitting and data transforming will generate two new matrices. One shows the topic distributions at each position (up right), and the other shows the probability of observing 6-mers from topics (down right).

**Figure 2.**
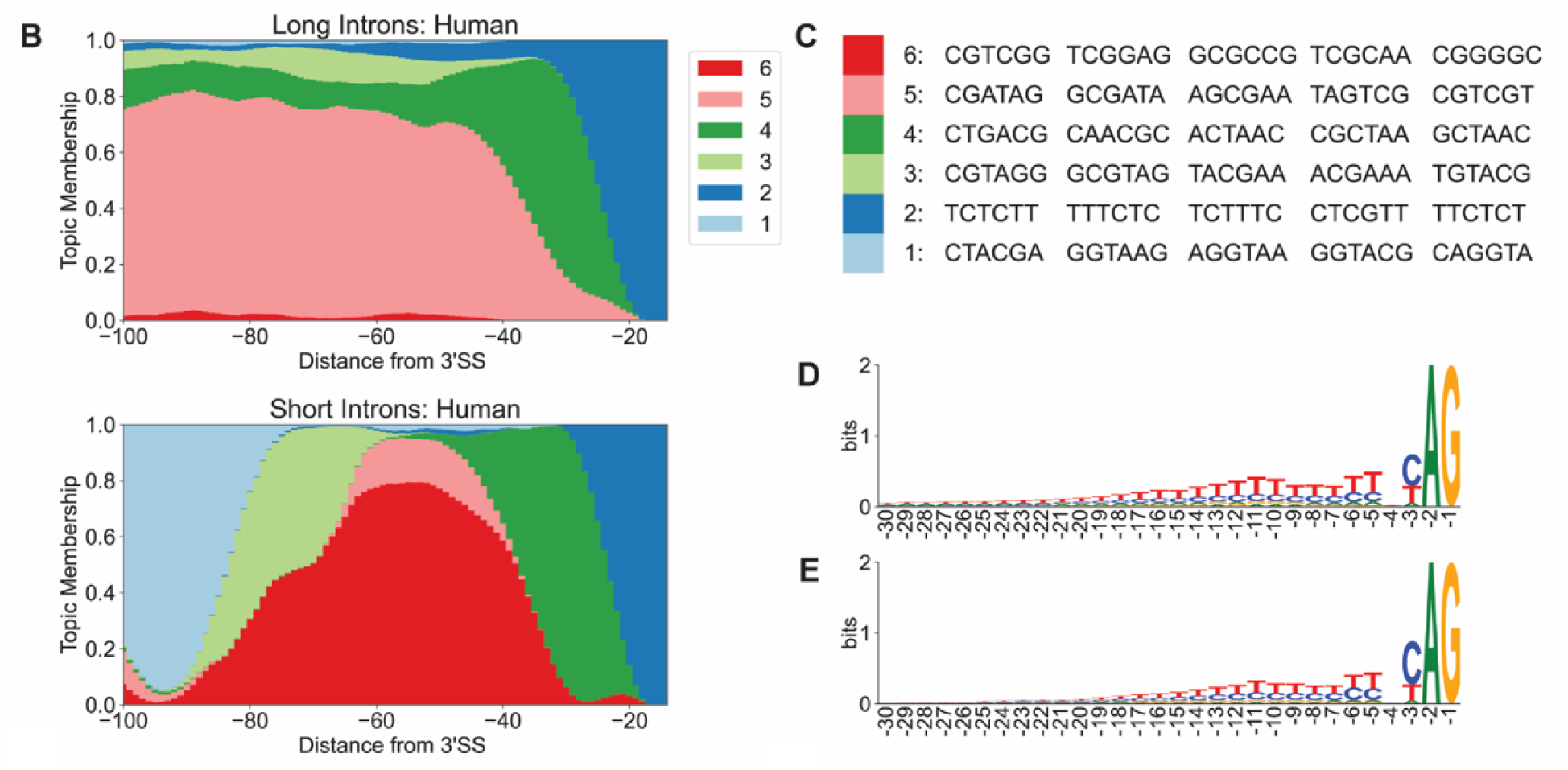
LDA characterization of short and long human introns using aligned positions as samples. B. Structure plots of long (n=10,000) and short (n=10,000) human introns. Hexamer features in 5 nt. windows starting at positions -100 to -13 upstream of 3’ splice sites (encompassing -100 to -3) were used as samples. C. Table of top 5 driving hexamer features in the six topics in Fig 2B. D. Sequence logo of 3’SS of long human introns. E. Sequence logo of 3’SS of short human introns.

**Figure 2.**
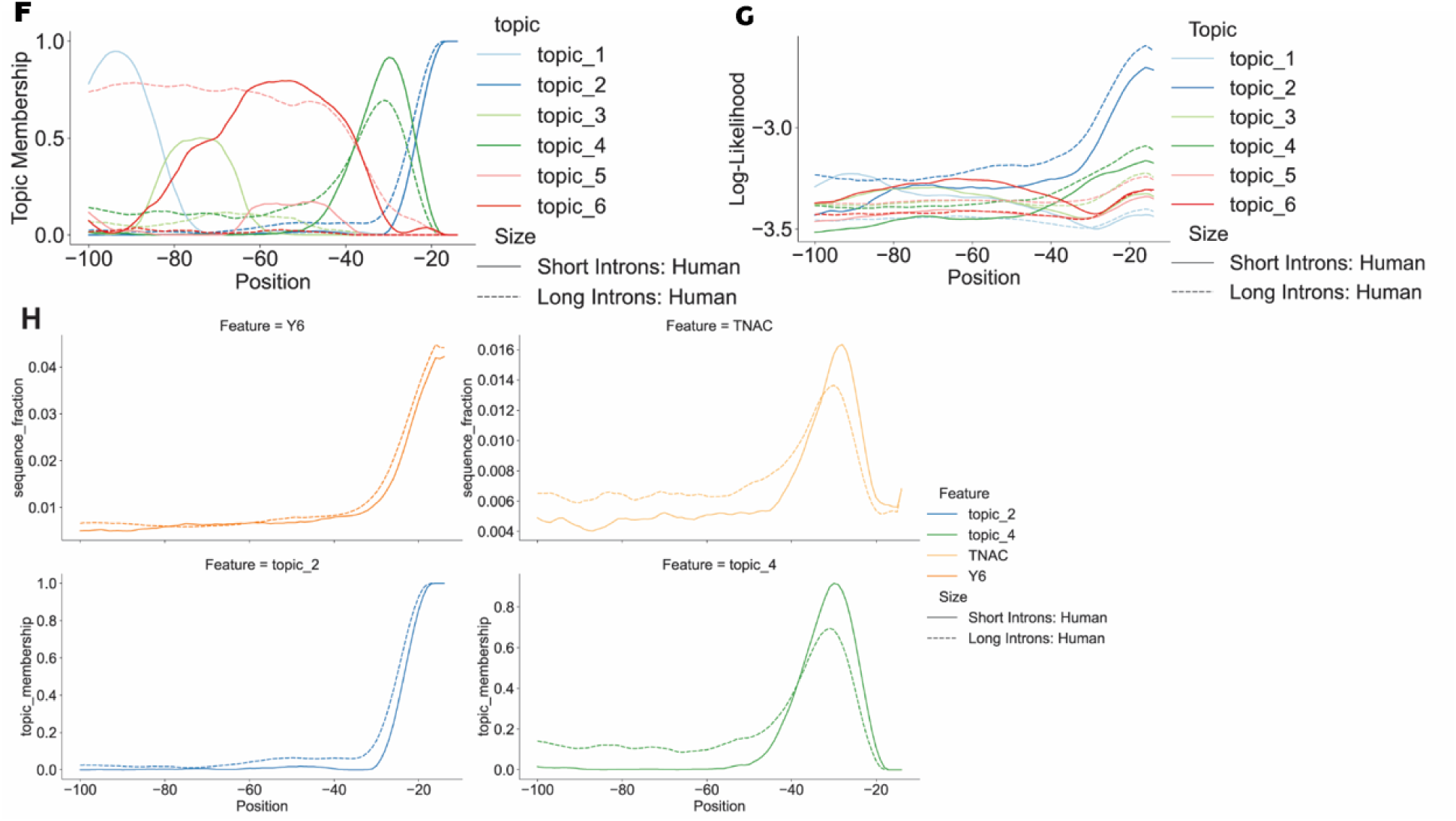
LDA characterization of short and long human introns using aligned positions as samples. F. Line plot of topic distribution across positions relative to the 3’SS of human introns. G. Line plot of the likelihood of observing the distribution of features at each position in human introns. H. Line plots of the branch site and pyrimidine tract consensus sequence signals and corresponding topic signals.

Both long and short human introns share a peak of topic 4 (branch site) membership at about -30 and a peak of topic 2 (pyrimidine tract) membership at about -15 relative to the annotated 3’ splice site (3’SS; Fig 2B). Thus, both the relative positions of topic 2 and topic 4 peaks and the interpreted topic meanings are consistent with the known sequence feature of human introns.

However, the fractional membership in topic 4 at some positions, such as -28 in short human introns, is close to 100%, much greater than the fraction of sequences with discernible branch site motifs. This is despite the difficulty of deriving a branch site consensus from sequence alone, and the absence of significant information content in position weight matrices aligned at the 3’SS (Fig 2D, 2E). This implies that topic 4 includes k-mers that are characteristic of the branch site region that do not match the branch site consensus, and may not be branch sites.

The distribution of topics from the model fitted with long and short introns shows the potential of LDA to identify functional motifs and distinguish sequence types. However, fractional representation, like that shown in Figure. 2B and 2F, does not indicate how well the topics describe the sequence, only which topics do best. To assess the performance of these topics in describing the raw sequences, we calculated the likelihood of reproducing k-mer compositions at every position by every topic (Figure 2G). The highest likelihood values are associated with known motifs (the branch site and pyrimidine tract) in the correct positions, which confirms the potential of unsupervised mixture models to identify functional signals. Also, the significant decay of the total log-likelihood as the distance from the 3’SS increases indicates that, as expected, these known motifs are much better described by the model than is bulk intron sequence far from the splice site.

The analysis also reveals differences between long and short introns. Compared with long introns, the region upstream of short human intron 3’ splice sites has more complex features, including an enrichment of topic 6 between 40 and 80 bp upstream; topic 6 is characterized by GC-rich driving k-mers and is almost entirely missing from long introns. Topic 3 is enriched between 65 and 85 bp upstream of the 3’SS, and topic 1 is enriched upstream of that. In contrast, topic 5, characterized hexamers containing AG or CG, is predominant in long introns. Because 48% of human intron sequences less than 150 bp are less than 100 bp (Figure S1H), the region shown in Fig. 2B includes the 5’SS’s, and most of the driving k-mers in topic 1 are subsets of the 5’ splice consensus sequence CAG|GTRAG (Figure. 2C).

By analyzing the Drosophila genome with the same pipeline, topics corresponding to the branch site and the polypyrimidine tract were again identified, and at the same positions as in the human sequence analysis; but with different driving k-mers (Figure S1A, S1B). As with human sequences, the relationship between these signals, and the difference between short and long introns, is not apparent in positional weight matrices alone (Figure S1C, S1D). Again, one of the significant differences between long and short introns is the presence of the 5’ splice site, represented here by topic 6, in the region modeled (Figure S1A, S1B). Again, all of the top 5 driving k-mers are subsets of the 5’ splice site consensus, AG|GTRAGT. This identifies topic 2 as a feature of exon sequences (Figure S1A, S1B).

To confirm that the position of the signals from mixture models is consistent with the known signals, we compared the location of the branch site and polypyrimidine tract motifs with the location of the corresponding topics. We used the fraction of 6-mers containing CUNA as a marker for the human branch site, and CUAA for the Drosophila branch site, and 6-mers with only pyrimidines for the pyrimidine tract. While only a very small fraction of sequences contain these specific motifs and nearly all sequences are represented by the corresponding topics, both signals have peaks at the same positions as the consensus signal curves, with similar differences between long and short introns (Figure 2H, S1G).

### Modeling sequences using positional k-mers distinguishes intron subtypes

To distinguish intron subtypes on the single sequence level, we applied mixture models using blocks of sequence around individual splice sites, instead of positions, as experimental units. The k-mer compositions of introns with different lengths are similar, which makes bulk k-mers unsuitable for this classification task (Figure S2A-D). However, mixture models indicated a difference between similar positions from introns of different lengths (Figure 2). Accordingly, we added positional information to the k-mer features, so that each feature is a k-mer at a specific position (e.g. CCAG_at_-4), and applied LDA to samples consisting of small packs of individual sequences (Figure 3A). We investigated the positional sequence features of 3’SS and 5’SS separately. To study the features of 3’SS from long (>=150bp) and short (<150bp) human introns, we selected 80 bp sequences that include 50bp from the intron and 30bp from the downstream exon. A model fitted with 60 packs of 250 single sequences (15,000 total), readily distinguished packs of 250 long or short introns (Figure 3B). Long intron sequence packs have topic 2 memberships around 90% while the short intron sequence packs have topic 2 memberships less than 15%. A separate set of sequences was used to test the model’s performance on single sequences, which generated a classification accuracy of 67.95% (Figure 3C, Methods).

**Figure 3.**
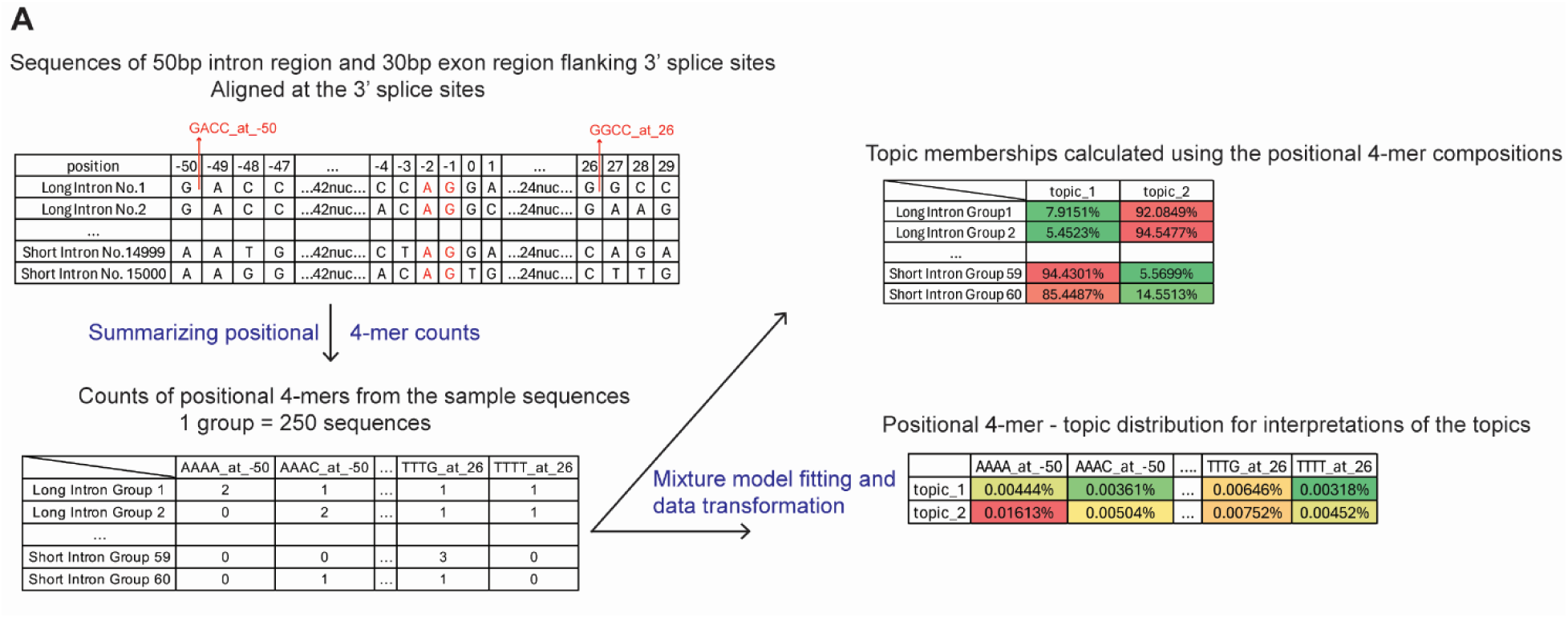
LDA characterization of short and long human 3’SS using individual sequences as samples. LDA was applied to samples corresponding to splice site regions from long and short introns from human (long>=150 nt.; short<150 nt.). A. Diagram of the LDA analysis of intron sequences aligned at 5’ or 3’ splice sites. In this diagram, we introduce the analysis of sequences flanking 3’ splice sites. First, sequences are aligned at the 3’ splice sites (up left), and the positions of A and G are -2 and -1. Then every 250 sequences are grouped into one sample group and the counts of positional 4-mers are summarized (down left). Examples of positional 4-mers are shown in the up left table highlighted in red color. LDA fitting and data transformation act as dimension reduction method and represent each sample as topic distributions, which can be interpreted using the topic-kmer table (right).

**Figure 3.**
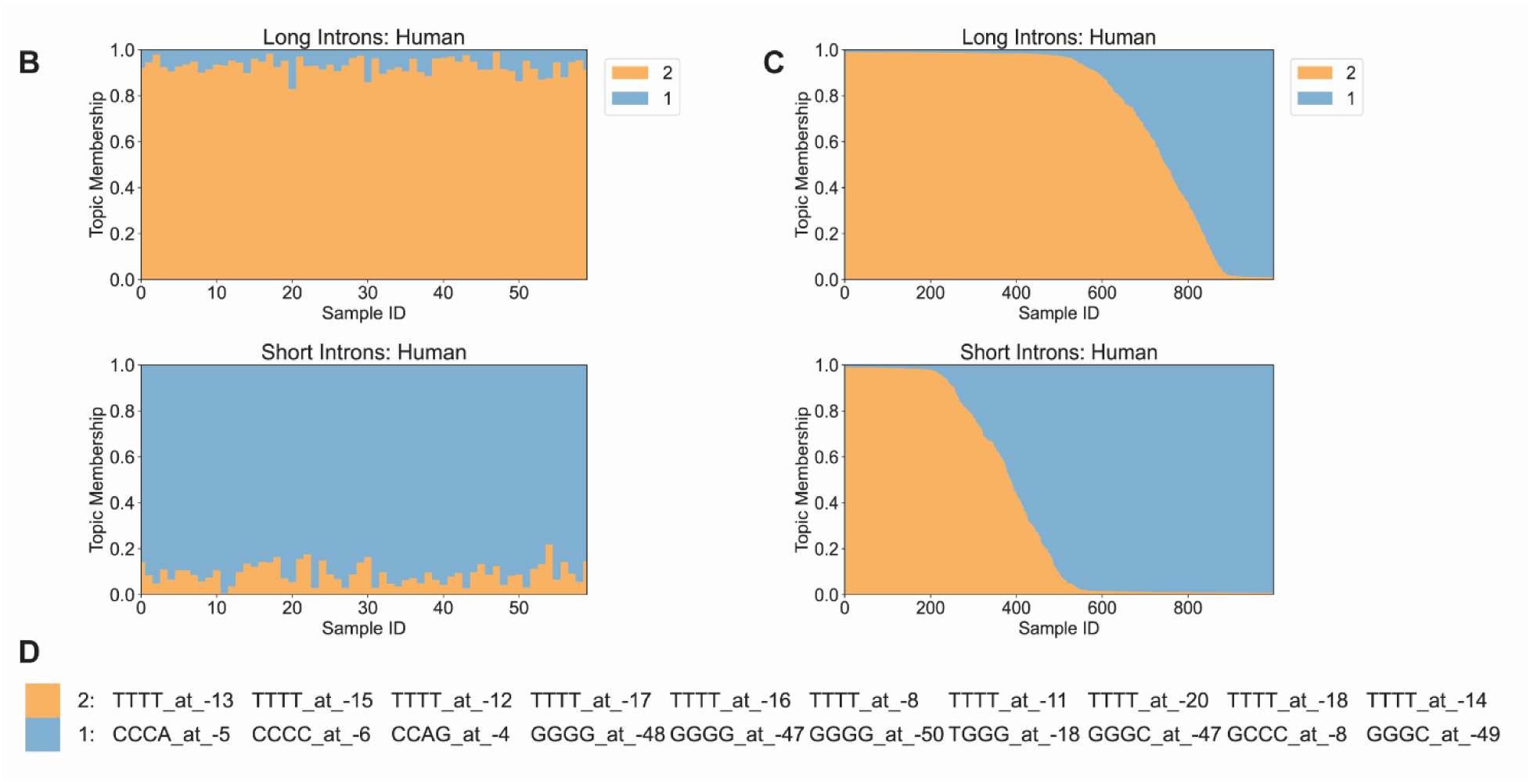
LDA characterization of short and long human 3’SS using individual sequences as samples. B. Structure plots of long (n=15,000) and short (n=15,000) human intron sequences (30 nt. of exon and 50 nt. of intron) near 3’SS. Every sample contains positional tetramer features from 250 sequences (15,000 sequences total). C. Structure plots of long (n=2,000) and short (n=2,000) human introns. Every sample is a single sequence. The topics are the same as the topics in Fig 3A. D. Top 10 driving positional tetramers from topics 1 and 2 from the analysis of 3’SS of human introns.

**Figure 3.**
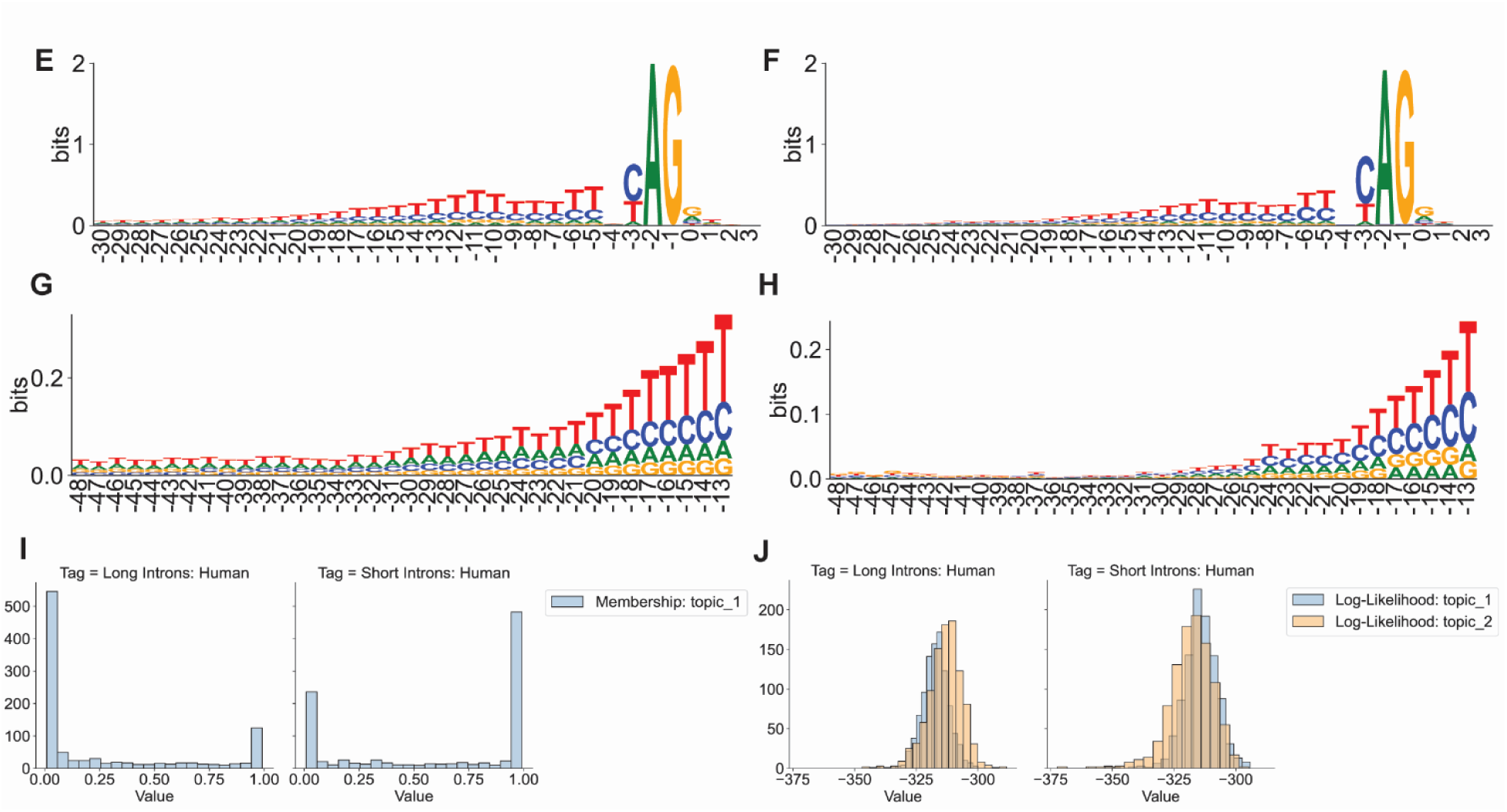
LDA characterization of short and long human 3’SS using individual sequences as samples. E. Sequence logo of the 3’SS from long human introns. F. Sequence logo of the 3’SS from short human introns. G. Sequence logo of positions -48 to -13 from long human introns. H. Sequence logo of positions -48 to -13 from short human introns. I. Distribution of topic 1 memberships of single sequences in Fig 3B. J. Distribution of the likelihood of generating sequences in Fig 3B by topics.

The same pipeline was applied to 80 bp sequences at the 5’SS, including 50bp from the intron and 30bp from the upstream exon (Figure S3), yielding a classification accuracy of 66.6%. Long and short introns differ in base composition in the region 17bp-20bp downstream of the 5’SS (Fig. S3F, S3G), and some driving k-mers reflect this (e.g. TTTT at 18 in topic 2 and GAGG at 22 in topic 1). However, the top-ranked driving k-mers are in the core 5’SS, including the invariant GT dinucleotide (Figure S3D, S3E). Based on the similarity between long and short intron consensus sequences, we conclude that LDA has the potential to find subtle differences between sequences.

Driving k-mers from these topics (Figure 3D) and sequence logos generated for the upstream region (Figure 3G, 3H) indicate that 1) the long intron group has a longer polypyrimidine tract, which is consistent with the observation of previous study (Yıldırım & Vogl, 2023); 2) topic 2, enriched in long introns, is characterized by runs of T in the pyrimidine tract (-20 to -5) while topic 1 is characterized by runs of C in the pyrimidine tract (-6 to -3); 3) topic 1 is characterized by GC-rich sequences upstream (-50 to -44). These differences characterize the most distinctive k-mers; there are many other differences. Though long introns have more sequences that are enriched in topic 2 and can be better represented by topic 2, both groups have many individual introns with significant memberships of the topics of the other labels (Figure 3C, 3I, 3J). It is important to bear in mind that the classification of introns by subtype may be more accurate than the assignment to size classes if mechanistically or biochemically distinct subtypes do not perfectly correspond to size classes.

The difference between intron subtypes was also revealed from Drosophila introns, with accuracies of 71.75% from the 5’SS and 66.6% from the 3’SS (Figure S4, S5). Similar to the human sequence analysis, intron sequences at 3’SS from the Drosophila genome also indicated a difference in the polypyrimidine tract regions. However, in Drosophila, the differences in the pyrimidine tract are more related to T-enrichment rather than the size of the signals. (Figure S4F, S4G).

### Modeling unaligned protein-coding sequences reveals features associated with reading frames and species

Many classic signals within genes are recurrent motifs over-represented in specific regions, such as the CpG islands in the promoter regions of human housekeeping genes. In addition, pervasive sequence differences characterize genomic regions, such as introns, and can be species-specific. To investigate the performance of mixture models in seeking over- and under-represented motifs from unaligned sequences, we analyzed the topic distribution of coding sequences in three reading frames with mixture models of 3 topics (Figure 4A).

**Figure 4.**
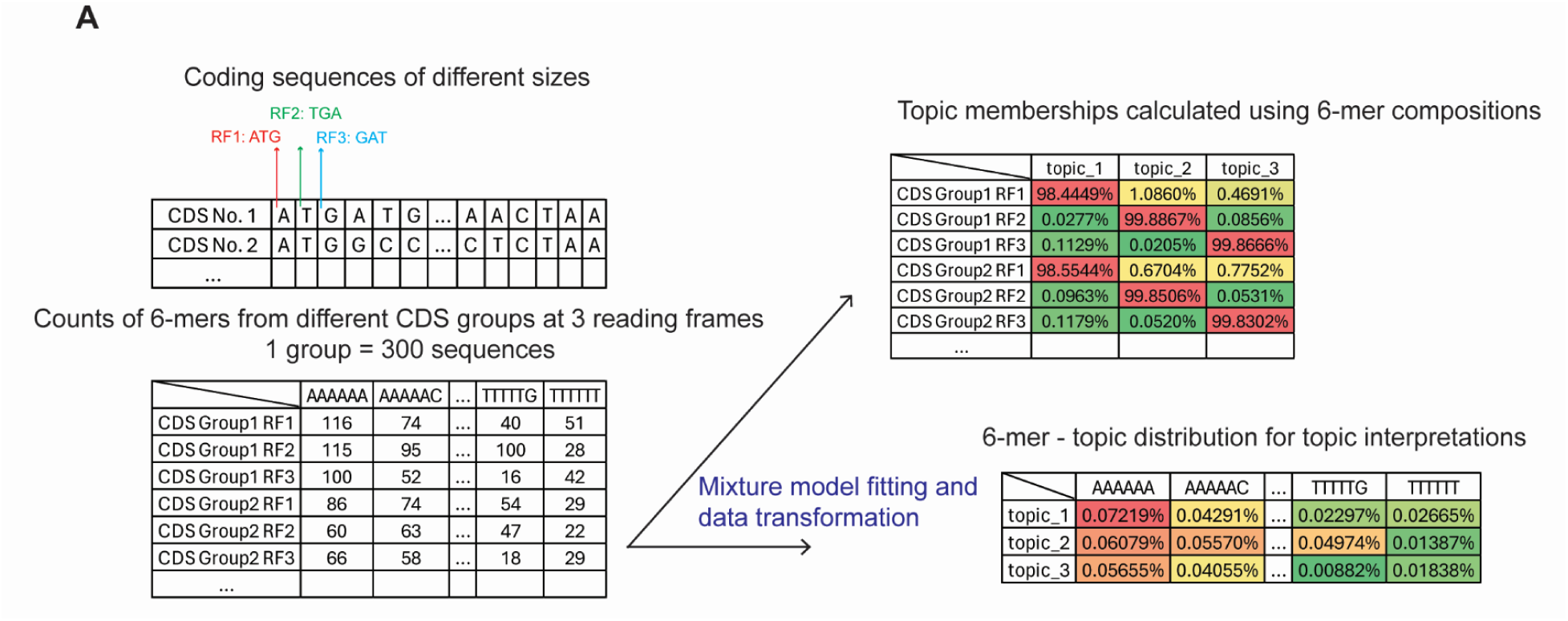
LDA characterization of reading frames using non-aligned coding sequences. LDA was applied to human CDS. A. Diagram of the LDA analysis of coding sequences. First, the reading frames from CDS are labeled (up left). From the example of the No.1 CDS, the 3-mer starting from the first base is from reading frame 1 and the 3-mer starting from the second base is from reading frame 2. Similarly, 3-mers starting from the fourth and fifth bases are from reading frames 1 and 2. Then every 300 CDS are grouped as a sample group, and the counts of 6-mers in each reading frame are summarized (down left). After LDA is fitted and the data is transformed, the sequence groups are represented as topic distributions with interpretable topics (right).

**Figure 4.**
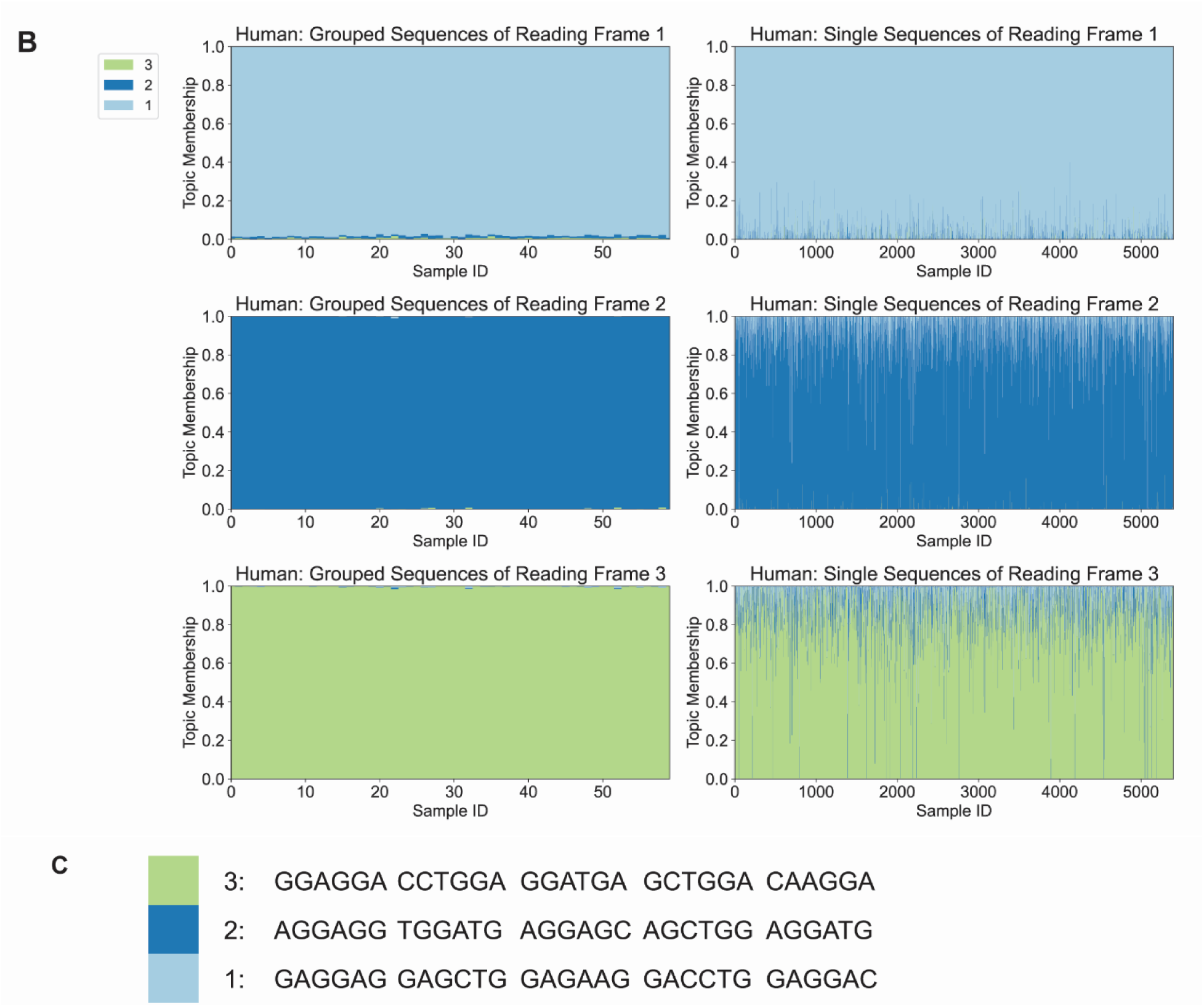
LDA characterization of reading frames using non-aligned coding sequences. LDA was applied to human CDS. B. Structure plots of human CDS sequences in reading frames 1, 2, and 3. The first column contains the model fitting data (18,000 CDS), in which each sample covers hexamer compositions of 300 sequences. The second column contains the topic distribution of single sequences (5,400 CDS). C. Top 5 driving hexamer features of 3 topics in Fig. 4B.

**Figure 4.**
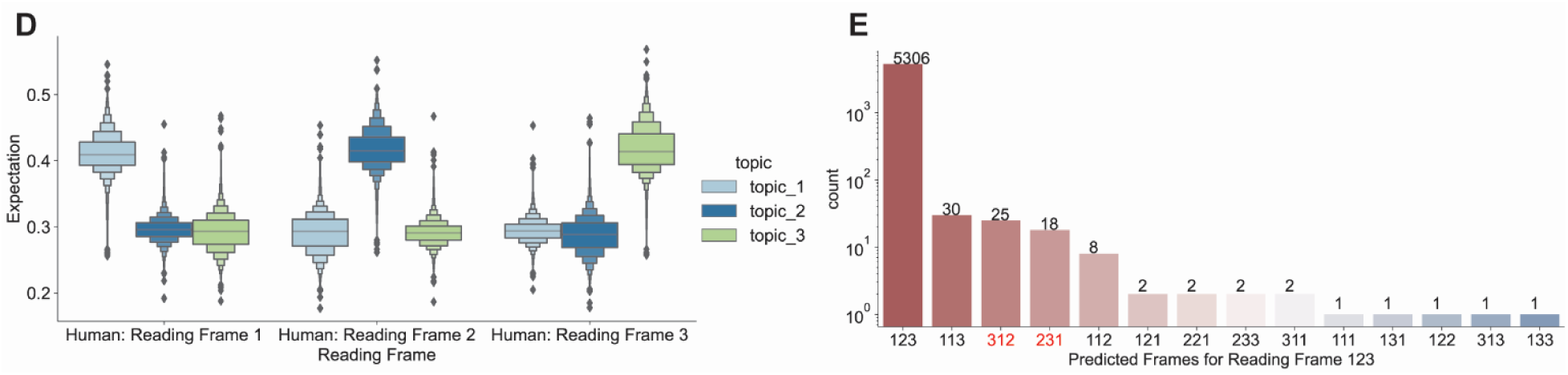
LDA characterization of reading frames using non-aligned coding sequences. LDA was applied to human CDS. D. Distribution of the expectation of the probabilities of generating each sample from the second column of Fig 4A by 3 topics. E. Barplot of the counts predicted reading frames of sequences in Fig S6A. The correct order is 123.

By randomly grouping 300 human CDS sequences as a single sample, we observed that each reading frame could be represented by a distinct topic, one with little distribution in other reading frames (Figure 4B, 4C). Since each topic is associated with one reading frame, we calculated the probability of generating each CDS using every topic. The expectation is consistent with the observation of the topic distributions (Figure 4D). The classification with a separate set of sequences shows an accuracy of 98.77% (Methods). However, some of the single sequences do show a misalignment of the topic and reading frame (Figure S6A). A comparison of the predicted and true labels showed that the predicted frames of more than 45% of the misclassified sequences were shifted by one, suggesting potential frame-shifting events or annotation errors (Figure 4E).

To test whether the model fitted by human sequences can be applied to other species, we transformed the Drosophila coding sequence k-mer composition using the same model. Topic distribution showed that the Drosophila reading frames follow the features identified from human sequences (Figure S6B, S6C), suggesting that the information learned by LDA may analyze sequences from novel species.

To explore if mixture models can distinguish the reading frames and species at the same time, we fitted a 6-topic model with both human and Drosophila CDS. The topic distribution suggests that each reading frame can be described by 2 topics, which have distinguishable distributions for different species (Figure 5A, 5B, 5C). We calculated the probability of generating each CDS using single topics to evaluate the association between topics and species. For all 3 pairs of topics from 3 reading frames, most sequences have higher expectations from topics associated with the source species (Figure 5D-F). Single sequence classification was performed, and 93.6% of the sequences were correctly predicted with regard to both the reading frame and species. Most of the misclassified sequences in this study were labeled as the wrong species (Figure S6D, S6E).

**Figure 5.**
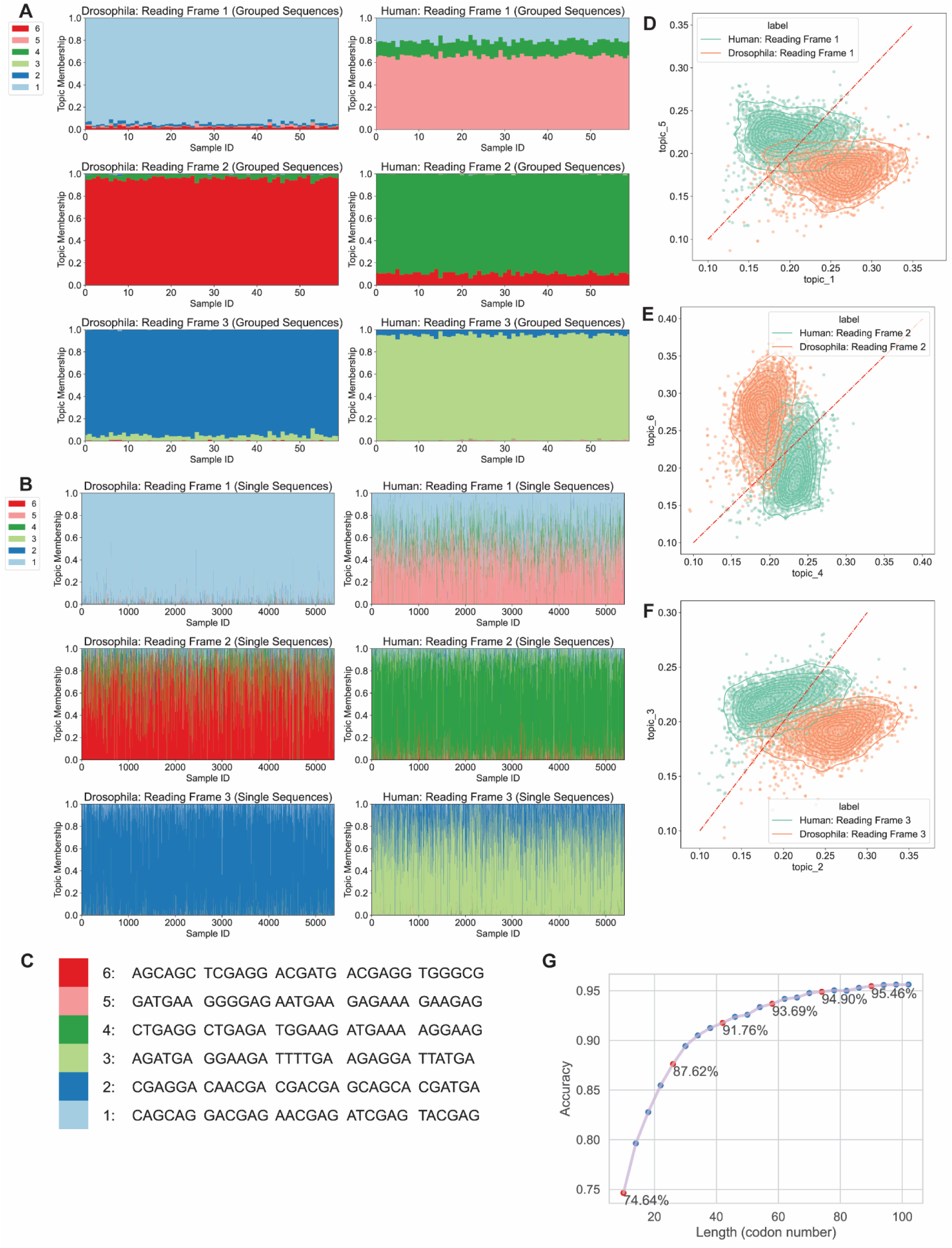
LDA characteristics of reading frames using non-aligned coding sequences. LDA was applied to both human and Drosophila CDS. A. Structure plots of human (18,000) and Drosophila (18,000) CDS sequences in reading frame 1, 2, and 3. Each sample contains the hexamers of 300 sequences. B. Structure plots of single sequences of human (5,400) and Drosophila (5,400) sequences. C. Top 5 enriched hexamer features of 6 topics in Fig 5A. D. The scatterplot of the probability expectations of generating reading frame 1 samples from Fig 5B using topic 1 or 5 from Fig 5C. E. The scatterplot of the probability expectations of generating reading frame 2 samples from Fig 5B using topic 4 or 6 from Fig 5C. F. The scatterplot of the probability expectations of generating reading frame 3 samples from Fig 5B using topic 2 or 3 from Fig 5C. G. Line plot of the accuracies of different lengths of subsequences of CDS to predict the reading frames.

If LDA were to be used to identify novel smORFs as coding sequences, it is useful to know how long a subsequence is necessary to identify the correct reading frame. To investigate this, we collected k-mer counts for short subsequences from 5,400 full coding ORFs in the CDS test set. These subsequences were between 10 and 100 codons drawn from the middle of the original coding sequences and were transformed using the LDA model fitted by full coding sequences. This method achieves an accuracy of 80% with only 18 codons, 90% with less than 34 codons and 95% with less than 74 codons (Figure 5G).

### Models based on coding regions identify potential protein-coding information in annotated UTRs and long non-coding RNAs (lncRNA)

To confirm that the signals mixture models identified are abundant and accurate, we analyzed the k-mer compositions from small open reading frames (smORFs) with at least 25 potential codons between the start and stop codons (Method) in human 5’UTR and 3’UTR. The topic distribution distinguished 5’UTR, 3’UTR, and CDS, but no differences were observed between the three frames of smORFs in the UTRs, where different samples share similar topic memberships (Figure S7A, S7B).

Often, 5’UTR smORFs can be translated into potentially functional protein products (Starck et al., 2016). The probability distribution of the UTR sequences showed that there are indeed some smORFs that have high expectations generated by the correct topics (Figure 6A, 6B). We analyzed single smORFs from 5’UTRs and recognized that 13,028 smORF CDS could be correctly classified into three reading frames using the classification model fitted by annotated CDS. Nevertheless, compared with real CDS, these UTR CDS showed different topic distributions of the correct topics associated with each reading frame (Figure 6A). To investigate the functions of genes with short 5’ UTR ORFs having correctly classified ORFs, we performed a Gene Ontology enrichment (Thomas et al., 2022) analysis. Most GO terms discovered are immune responses, sensory perceptions, and metabolic processes (Figure S7C).

**Figure 6.**
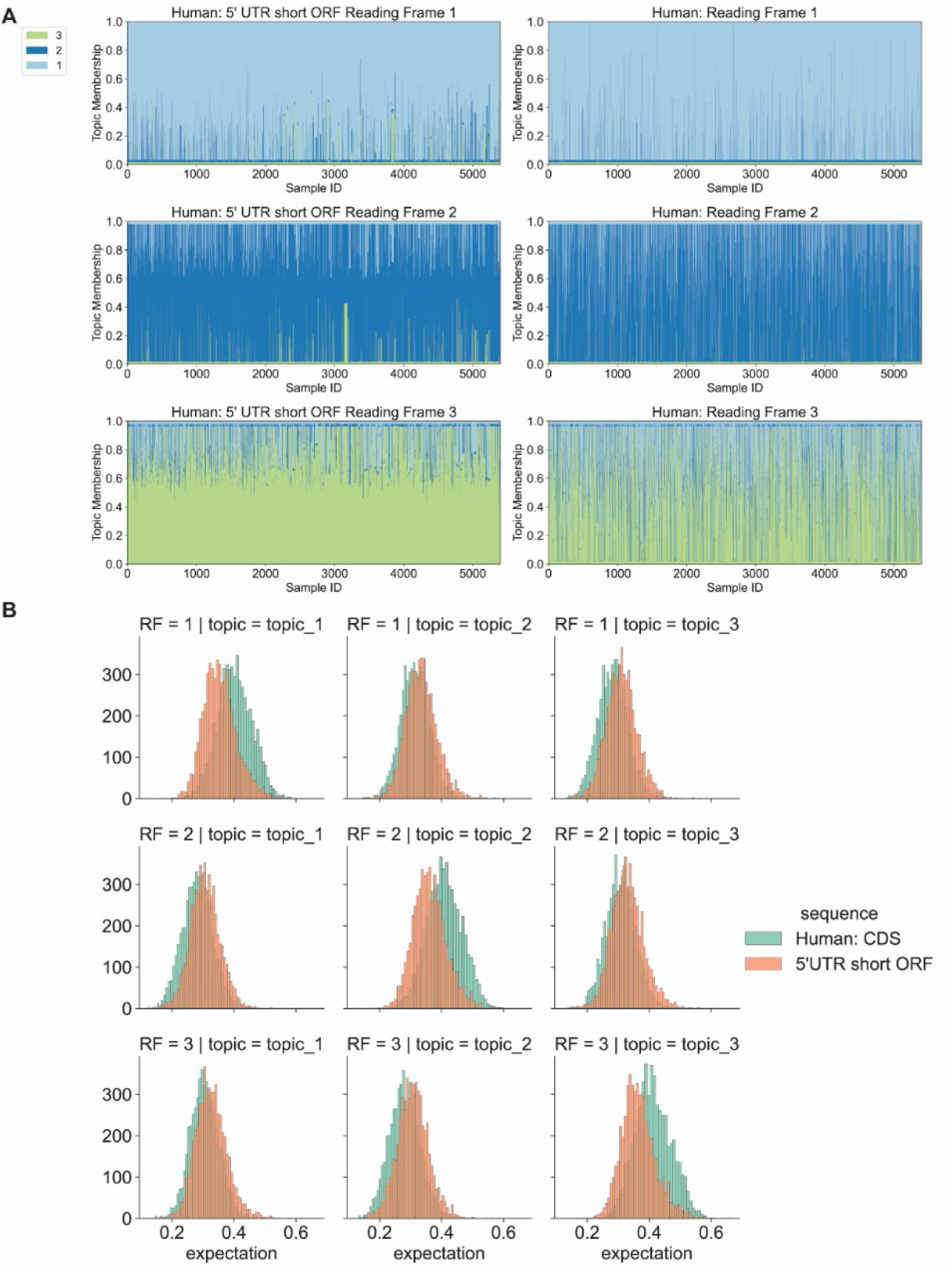
Identification of potential coding reading frames in human UTRs. The LDA model fitted by annotated CDS in Fig 4A was applied to small reading frames from human UTRs, each of which had a pair of start and stop codons and at least 25 potential codons. A. Structure plots of single sequences from human small reading frames in UTR5 and reading frames from CDS. Data were transformed by the model fitted in Fig 4A. 5,400 of UTR5 sequences that follow the CDS topic distribution patterns were selected to show. B. Distribution of the probability expectations of generating small reading frame samples from UTR5 and reading frames from human CDS using topics fitted in Fig 4A.

We also analyzed human lncRNA sequences using the same method. The distribution of sequence-generating expectations by topics of lncRNA also indicates that 10,532 sequences have similar features of coding sequences. Compared with UTRs, lncRNA showed patterns of topic distribution more similar to CDS. After reading frame prediction, we identified several lncRNAs with CDS features (Figure 7A).

**Figure 7.**
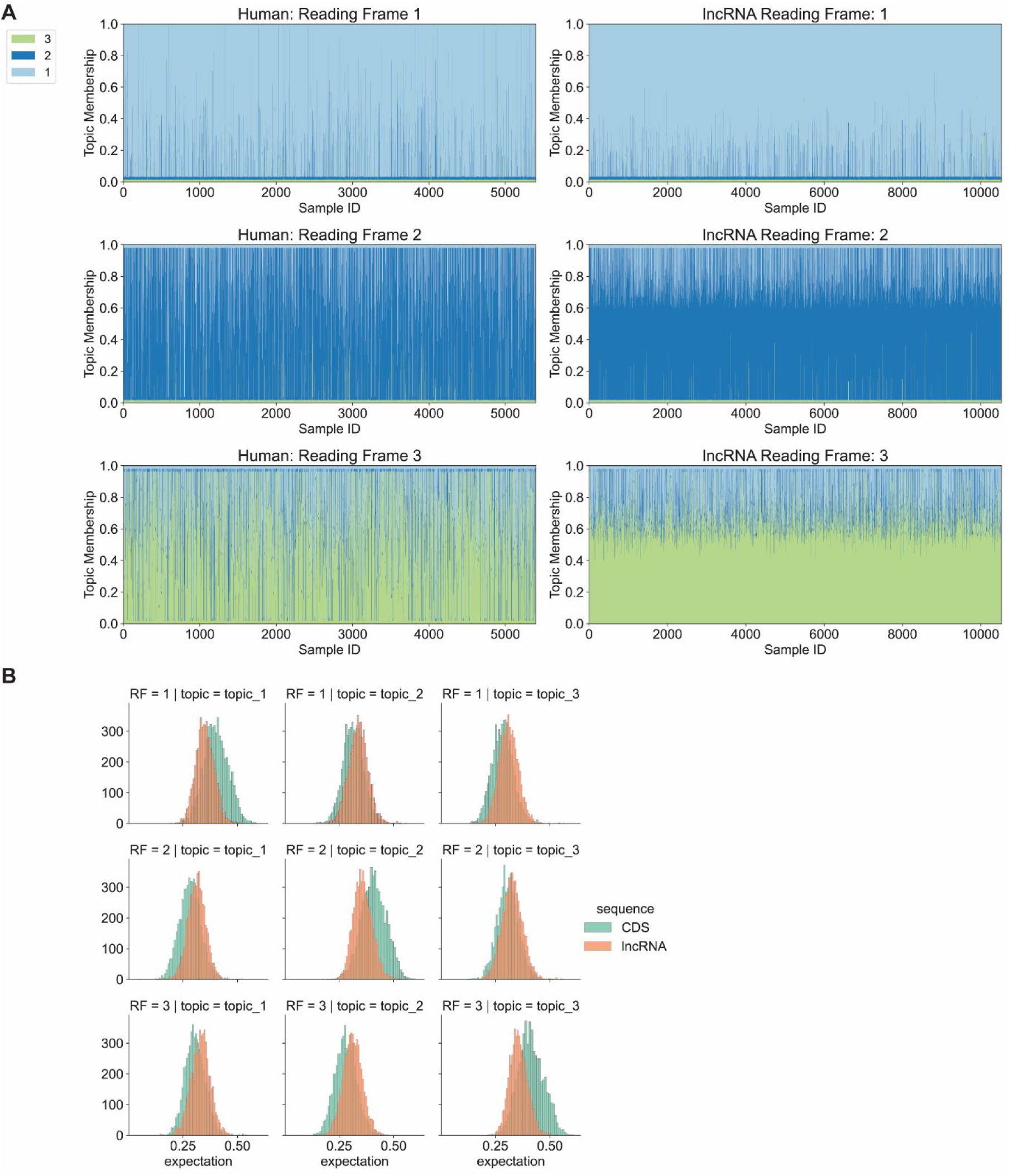
Identification of potential coding reading frames in human lncRNAs. The LDA model fitted by annotated CDS in Fig 4A was applied to small reading frames from human lncRNAs, each of which had a pair of start and stop codons and at least 25 potential codons. A. Structure plots of single sequences from human small reading frames in lncRNA and reading frames from CDS. Data were transformed by the model fitted in Fig 4A. 5,400 of lncRNA sequences that follow the CDS topic distribution patterns were selected to show. B. Distribution of the probability expectations of generating small reading frame samples from lncRNAs and reading frames from human CDS using topics fitted in Fig 4A.

## Discussion

Here we have shown the utility of mixture models using LDA for visualizing and interpreting signals in nucleotide sequences. LDA readily identifies topics that correspond to sequence subtypes, and reports the proportional membership of those topics in each sequence. Because topics are interpretable, it is possible to easily identify functional motifs that function in biological processes. Examples here include the branch site consensus (topic 2 in Figure 2C: RCTRAC), the 5’ splice site (topic 3 in Figure 2C: AGGTA), and an unknown G-rich motif characteristic of short human introns (topic 6 in Figure 2C). In addition, LDA can uncover subtypes of sequences based on differences in topic distribution. Examples include subtypes of intron correlated with length, reading frame and species. Finally, the expectation that a sequence is derived from a topic can be used as a score to identify sequences with potential functions, such as potentially coding small ORFs in 5’ UTRs.

### Motif discovery

Because topics are interpretable, it is possible to identify motifs using LDA. A motif can be readily interpreted by viewing the enriched or driving k-mers from the enriched topics. While the basic probabilities of the observations for each topic could be used, top k-mers based on raw probabilities can be shared between topics, and alternative methods to identify the driving features are preferable. We identified distinctively distributed (“driving”) k-mers using K-L divergence (following Dey et al., 2017). It is the application of this method (Fig. 2C) that yielded driving k-mers drawn from known consensus sequences for the branch site, 5’ splice site and pyrimidine tract, as well as an unfamiliar G-rich motif characteristic of short introns. Driving features that are k-mers with sequence overlap can be used to identify motifs that can be represented by sequence logos.

### Sequence subtypes

LDA identifies sequence subtypes directly from topic membership, whether based on bulk k-mer composition or positional k-mer counts. However, subtype recognition can be improved in several ways. In this paper we have used sequences aligned at a splice site, or by reading frame. This creates a contrast in k-mer frequencies between samples based on that alignment. It is necessary to use positional k-mers to identify subclasses of sequence that have not been identified in advance. Sequences of variable length that are not readily aligned may be more amenable to bulk k-mer compositions.

Pooling sequences prior to model fitting is useful to overcome the problem that single sequences have 0 occurrences of many features, which is especially true for positional k-mer values. Thus, we improved model performances by randomly grouping sequences from the same known group as one single sample. This approach increased the distinction between known sequence groups. Single sequences transformed by the model fitted by grouped sequences have much higher variance in the topic distributions. We believe that grouping sequences can magnify the impact of enriched k-mers and push the model to generate topics to describe the generative process of two groups instead of single sequences. However, the enriched features in a group may not occur in every single sequence. One potential flaw of this method is that sequences with abundant repetitive patterns can introduce bias, potentially reducing the signal of true subtle features.

### Overview, caveats and prospects

Here we have used LDA and k-mer features to classify and characterize nucleotide sequences. One obvious extension of this work would be application of LDA to protein sequences. Another would be to include features, such as accessibility in chromatin, that are not properties of the sequence *per se*.

As a generative statistical model, LDA derives topics that describe the data used to fit the model. These topics can then be used to score novel sequences using the expectation or likelihood that those novel sequences were generated from the topic. In this way, novel sequences that potentially share biological functions with the sequences in the fitted set sharing those topics can be identified. However, topics that are not observed by the model cannot be predicted.

We recommend LDA as a tool that will be broadly useful for describing the properties of sequences in a functional class.

## Methods

### Overview of sequence feature analysis with LDA

For each analysis, matrices of k-mer feature counts were generated for each sample (see below). The latent Dirichlet allocation (LDA) algorithm was implemented using the sci-kit learn (Pedregosa et al., 2011) package. Driving k-mers from each topic are pulled by the calculation of distinctiveness through the Kullback-Leibler divergence (Dey et al., 2017). Structureplot visualization (Pritchard et al., 2000) is implemented on Matplotlib package (Hunter, 2007). Sequence logos were generated with the Logomaker package (Tareen & Kinney, 2020). Matrix computations are performed with Numpy (Harris et al., 2020) and Pandas (McKinney, 2010).

### Analysis using positions as samples

To evaluate intron sequence features associated with positions, we randomly selected human (hg19) and Drosophila (dm6) introns and aligned them at the annotated 3’SS (O’Leary et al., 2016). Introns were labeled as long or short if they were greater or less than 150 bp (human) or 80 bp (Drosophila). Feature k-mer length, window size, pack size (the number of individual sequences per sample) and the number of topics are all options. Here, we counted 6-mers starting at every position from 100bp to 7bp upstream of 3’SS, so that the region from -100 to -2 was represented. A sliding window of length 5 was applied to batch the counts at adjacent positions together. LDA was applied to fit 6 topics.

### Branch site and polypyrimidine tract signal evaluations

We calculated the signal of the branch site by counting the fractions of k-mers that match the human (TNAC) and Drosophila (TAAC) branch site consensus. The polypyrimidine tract signals were evaluated by counting the fractions of k-mers with only Cs and Ts.

### Positional k-mer counts as features

Positional features incorporate both sequence and position in the alignment. For example, the sequence AGTTAT, starting at position 1, would have AGTT_1, GTTA_2 and TTAT_3 as features. For both human (hg19) and Drosophila (dm6) splice sites from long and short introns, we randomly selected 16,000 sequences comprising 30 exon and 50 intron nucleotides. 15,000 sequences were used for model fitting, and the remaining 1,000 sequences for model testing. Sequence feature analysis for 3’SS and 5’SS were performed separately for each species.

### Model fitting by pooling sequences

Larger samples were obtained by pooling multiple sequences of the same type. For example, in the analysis of long and short introns, we used 60 samples from 15,000 introns (250 sequences per sample) to fit the model with 2 topics.

### Scoring a sequence’s fit to a topic

The intron subtype analysis was applied with 2 topic-LDA model. We hypothesized that the 2 topics generated are associated with long and short introns. By calculating the likelihood of generating each sequence from a topic, we were able to evaluate the information obtained from the sequences.

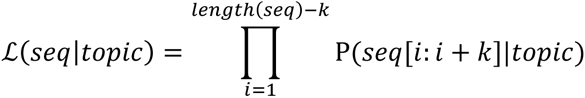

### K-mer composition as features and application to coding sequences

To evaluate the performance of LDA in differentiating sequence subtypes, we utilized protein-coding sequences (CDS) of human (hg19, CCDs) and Drosophila (dm6). For each sequence, we counted the occurrences of 6-mers in 3 reading frames. We used 60 samples from 15,000 CDS (300 sequences per sample) to fit the model with 6 topics.

To evaluate the ability of this model to identify the correct reading frame, we randomly picked 5,400 sequences that were not used in the model fitting step to test the performance on single sequences. Topics generated from the model fitting step were used to transform single sequence k-mer counts into topic memberships.

### Sequence subtype prediction

We implemented a simple classifier for predicting sequence labels in two steps, associating topics with labels and calculating sequence scores for each label. To associate topics with labels, we built a scoring system. For a given label, the score from a topic is the log2 of the fraction between the average membership in samples from such label and the overall average membership.

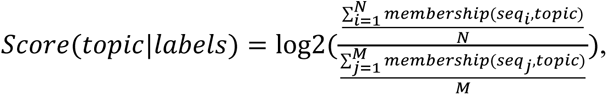

where N is the number of sequences of the label of interest and M is the total number of sequences in the analysis.

When calculating the result of a sequence, we multiplied the sample-topic matrix with the topic-score matrix and picked the highest score as the prediction.

### Frame-shift analysis

From the single sequences in the test set, we used the classifier to predict the reading frames based on each sequence’s k-mer counts from 3 frames. If the predicted reading frames from the original counts of reading frames 1, 2, 3 were 2, 3, 1 or 3, 1, 2, the sequence would be labeled as a potential frame-shift event.

### Prediction of protein coding in human UTR regions and lncRNAs

We scanned 5’ UTR and 3’ UTR sequences for a pair of start codon and stop codon in the same frame with at least 25 potential codons between them. The subsequence of the UTR from the start codon to the stop codon was identified as a small ORF reading frame 1 sequence, which was utilized to generate k-mer counts of 3 small reading frames. We predicted the reading frames of the UTR sequences and picked sequences with the correct prediction output.

### Prediction of subsequences

Subsequences from the human CDS test set (n = 5,400) were extracted by fixing the center of each sequence and expanding upstream and downstream equally to obtain coding subsequences centered on the middle of the original ORF. These subsequences were then transformed into topic memberships using the LDA model fitted with the full-length CDS test set.

All code is available on GitHub repository (github.com/xnao25/XMUSE)

---

**Figure S1.**
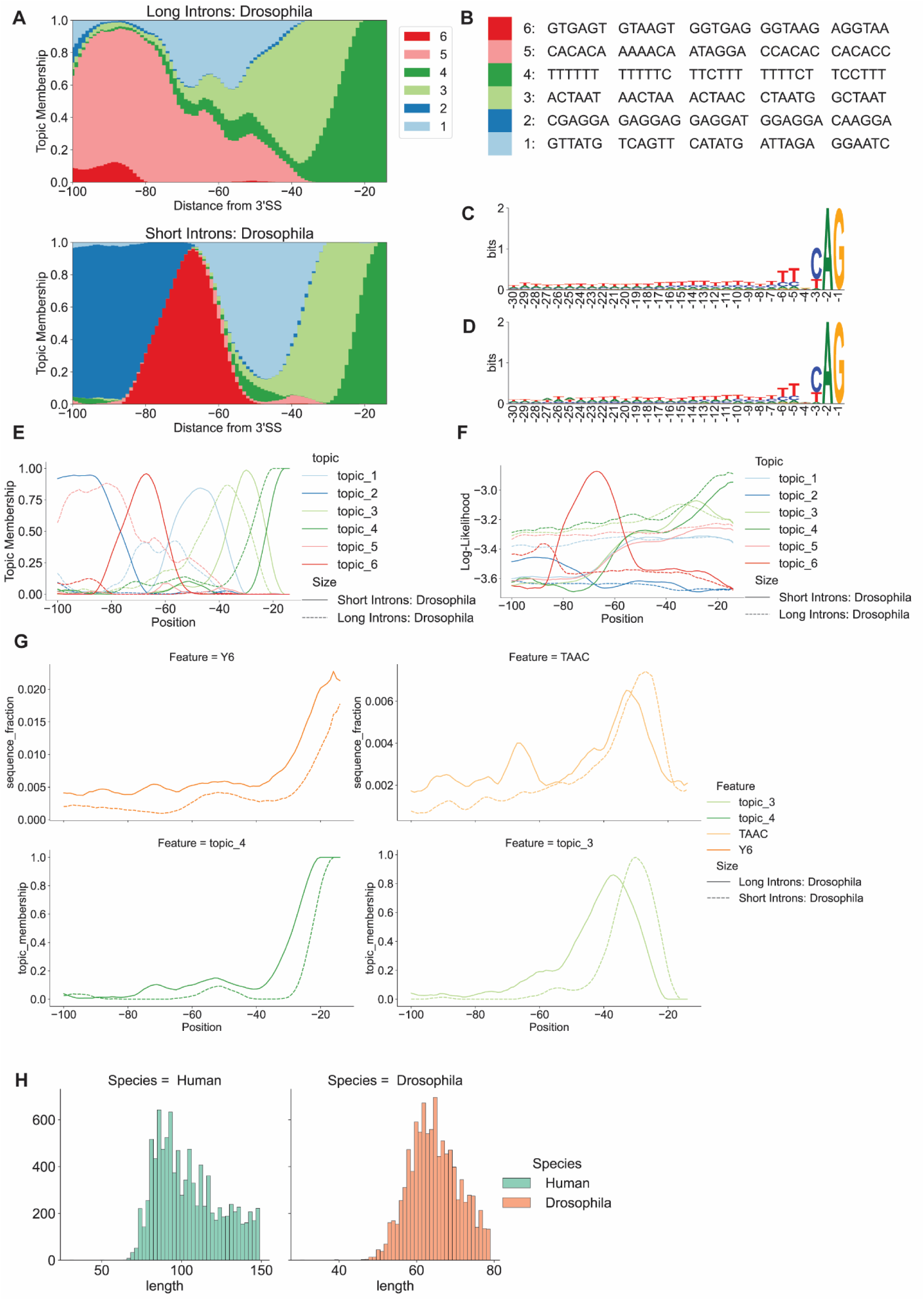
LDA characterization of short and long Drosophila introns using aligned positions as samples. LDA was applied to samples corresponding to positions in an alignment of 3’ splice sites from long and short introns from Drosophila (long>=80 nt.; short<80 nt.). A. Structure plots of long (n=10,000) and short (n=10,000) Drosophila introns. Hexamer features starting at positions -100 to -13 upstream of 3’ splice sites (encompassing -100 to -3) were used as samples. B. Table of top 5 enriched hexamer features in the six topics in D. C. Sequence logo of 3’SS of long Drosophila introns. D. Sequence logo of 3’SS of short Drosophila introns. E. Line plot of topic distribution across positions relative to the 3’SS of Drosophila introns. F. Line plot of the likelihood of observing the distribution of features at each position in Drosophila introns. G. Line plots of the branch site and pyrimidine tract consensus sequence signals and corresponding topic signals. H. The distribution of short intron sizes in the analysis of Fig 2 and Fig S1.

**Figure S2.**
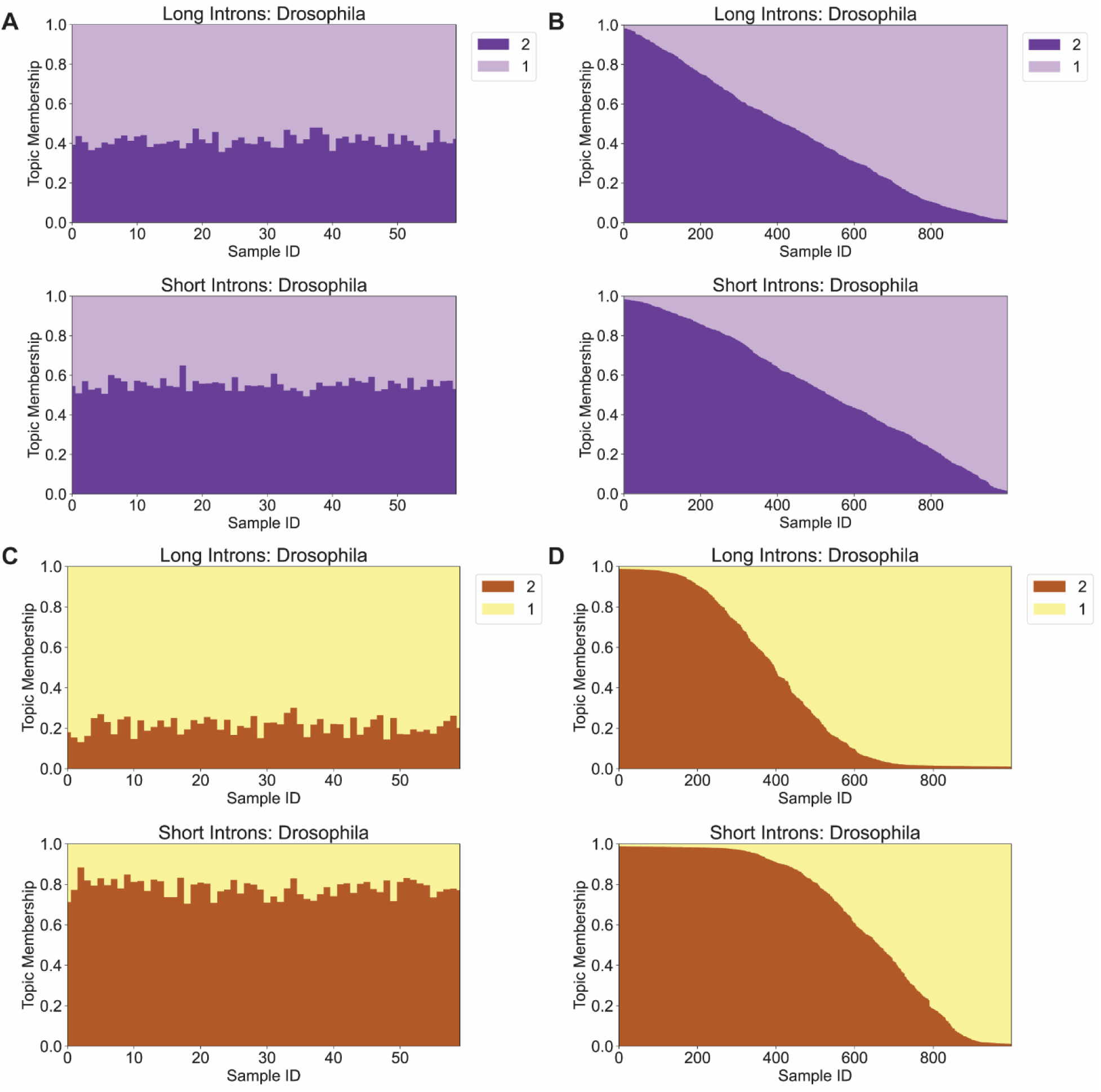
Non-positional feature analysis can not distinguish intron subtypes from Drosophila introns. LDA was applied to Drosophila intron sequences to identify differences between long and short introns using single sequences. Unlike the result in Fig S1, non-positional features are not able to describe the differences between the two groups. A. Structure plots of long (n=15,000) and short (n=15,000) Drosophila intron sequences (30 nt. of exon and 50 nt. of intron) near 3’SS. Every sample contains tetramer features from 250 sequences (15,000 sequences total). B. Structure plots of long (n=2,000) and short (n=2,000) Drosophila intron sequences near 3’SS. Every sample is a single sequence. The topics are the same as the topics in Fig S2A. C. Structure plots of long (n=15,000) and short (n=15,000) Drosophila intron sequences (30 nt. Of exon and 50 nt. Of intron) near 5’SS. Every sample contains tetramer features from 250 sequences (15,000 sequences total). D. Structure plots of long (n=2,000) and short (n=2,000) Drosophila introns sequences near 5’SS. Every sample is a single sequence. The topics are the same as the topics in Fig S2C.

**Figure S3.**
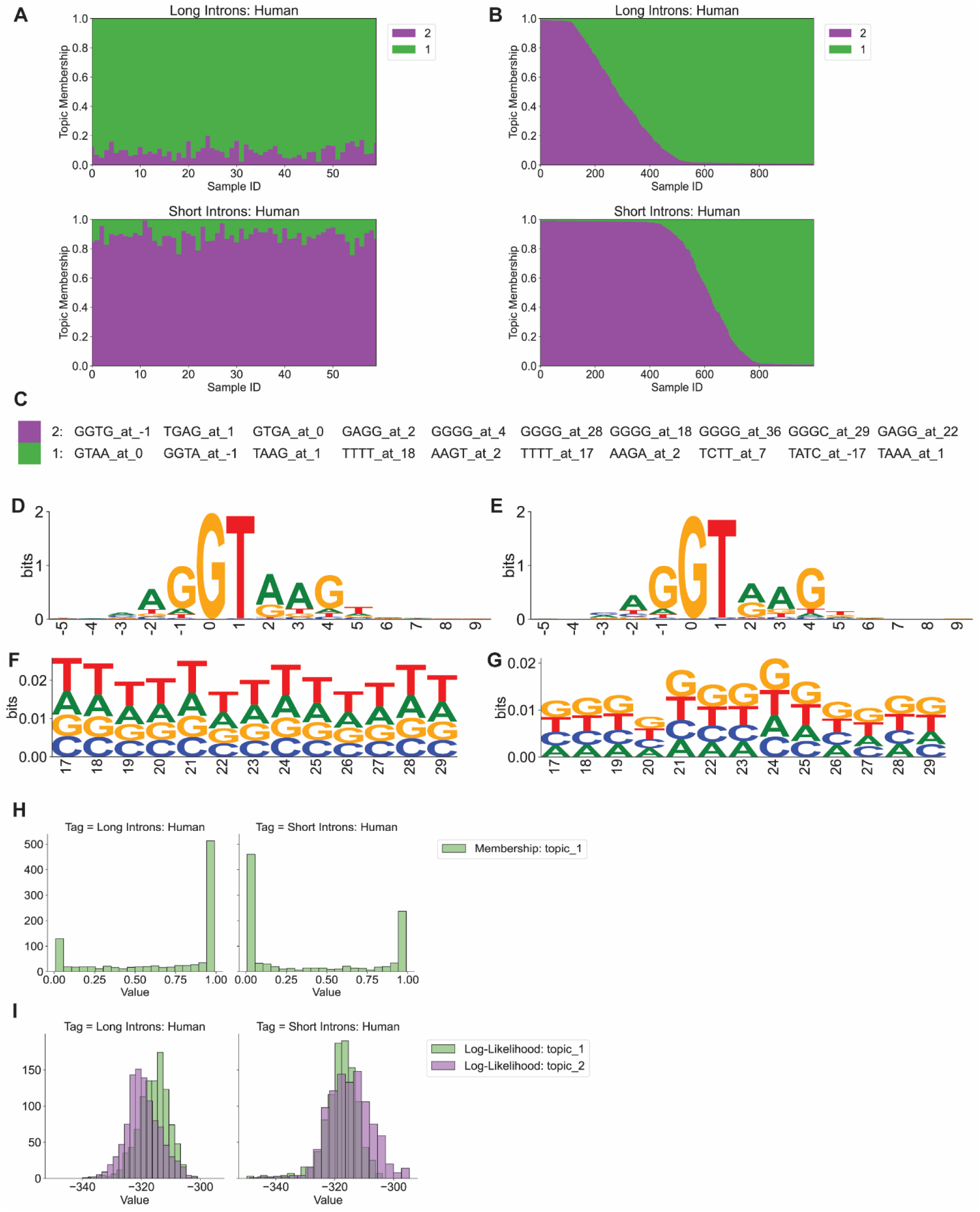
LDA characterization of short and long human 5’SS using individual sequences as samples. LDA was applied to samples corresponding to splice site regions from long and short introns from human (long>=150 nt.; short<150 nt.). A. Structure plots of long (n=15,000) and short (n=15,000) human intron sequences (30 nt. of exon and 50 nt. of intron) near 5’SS. Every sample contains positional tetramer features from 250 sequences (15,000 sequences total). B. Structure plots of long (n=2,000) and short (n=2,000) human introns. Every sample is a single sequence. The topics are the same as the topics in Fig S3A. C. Top 10 enriched positional tetramers from topics 1 and 2 from the analysis of 5’SS of human introns. D. Sequence logo of the 5’SS from long human introns. E. Sequence logo of the 5’SS from short human introns. F. Sequence logo of positions 17 to 29 from long human introns. G. Sequence logo of positions 17 to 29 from short human introns. H. Distribution of topic 1 memberships of single sequences in Fig S3B. I. Distribution of the likelihood of generating sequences in Fig S3B by topics.

**Figure S4.**
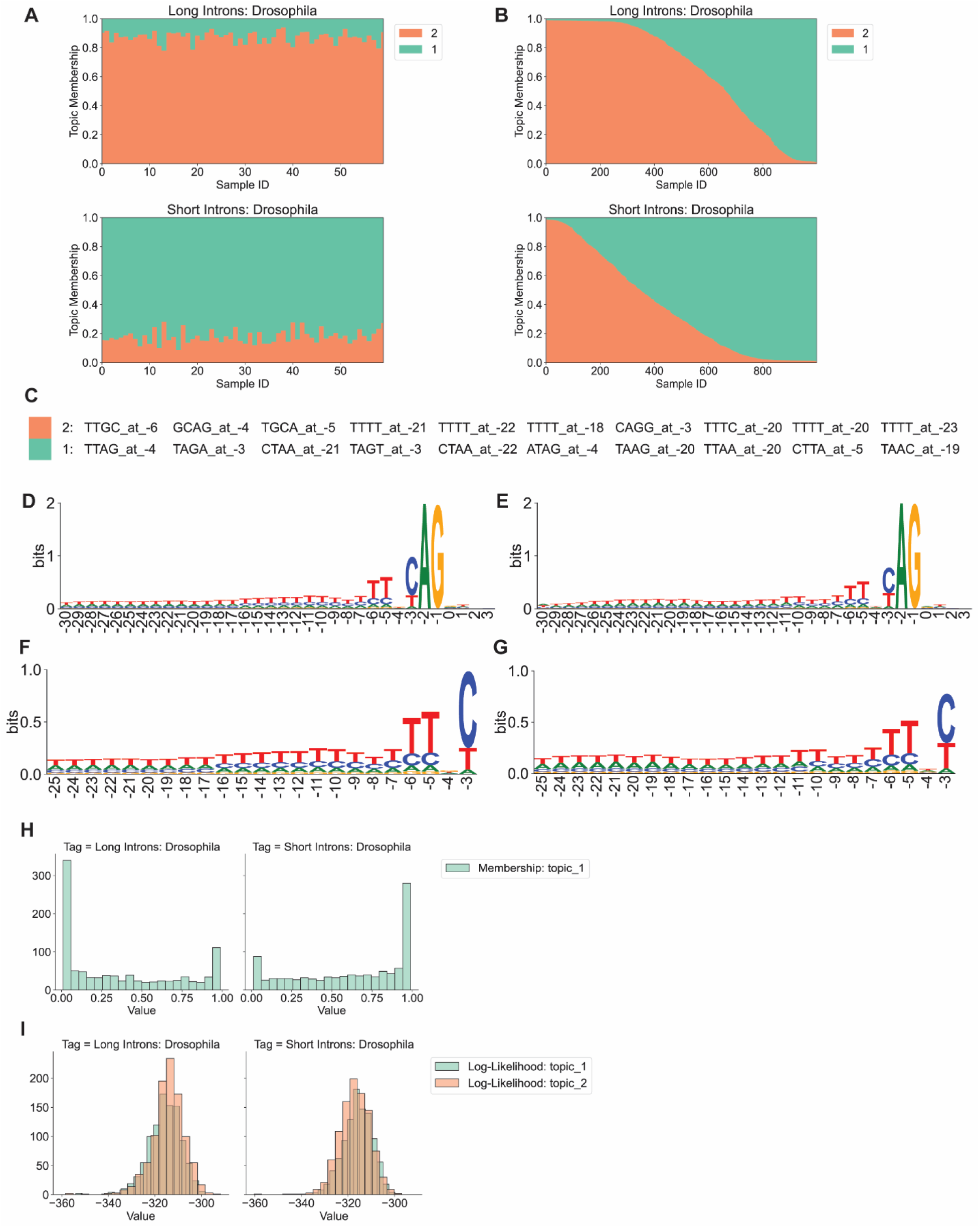
LDA characterization of short and long Drosophila 3’SS using individual sequences as samples. LDA was applied to samples corresponding to splice site regions from long and short introns from Drosophila (long>=80 nt.; short<80 nt.). A. Structure plots of long (n=15,000) and short (n=15,000) Drosophila intron sequences (30 nt. of exon and 50 nt. of intron) near 3’SS. Every sample contains positional tetramer features from 250 sequences (15,000 sequences total). B. Structure plots of long (n=2,000) and short (n=2,000) Drosophila introns. Every sample is a single sequence. The topics are the same as the topics in Fig S4A. C. Top 10 enriched positional tetramers from topics 1 and 2 from the analysis of 3’SS of Drosophila introns. D. Sequence logo of the 3’SS from long Drosophila introns. E. Sequence logo of the 3’SS from short Drosophila introns. F. Sequence logo of positions -25 to -3 from long Drosophila introns. G. Sequence logo of positions -25 to -3 from short Drosophila introns. H. Distribution of topic 1 memberships of single sequences in Fig S4B. I. Distribution of the likelihood of generating sequences in Fig S4B by topics.

**Figure S5.**
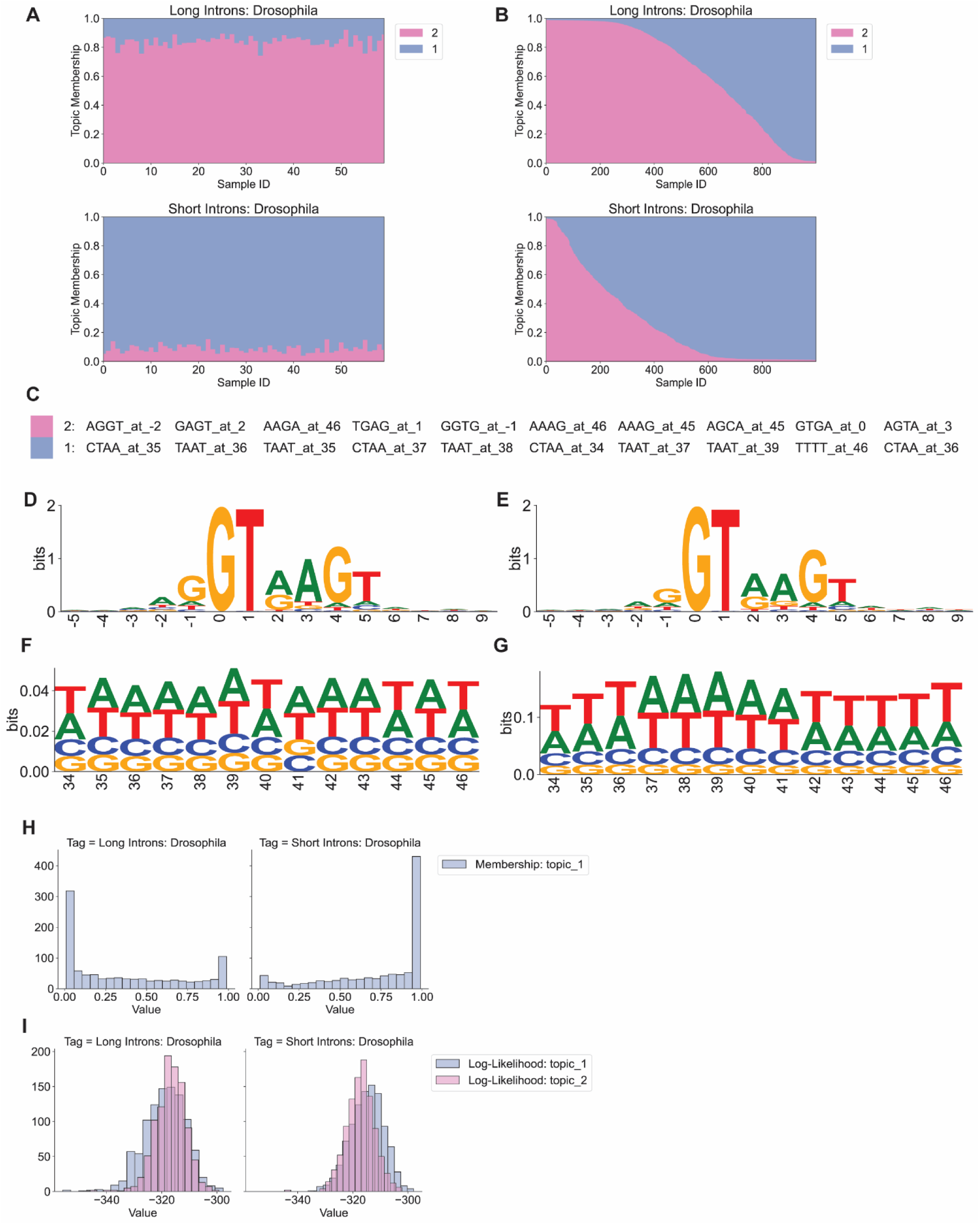
LDA characterization of short and long Drosophila 5’SS using individual sequences as samples. LDA was applied to samples corresponding to splice site regions from long and short introns from Drosophila (long>=80 nt.; short<80 nt.). A. Structure plots of long (n=15,000) and short (n=15,000) Drosophila intron sequences (30 nt. of exon and 50 nt. of intron) near 5’SS. Every sample contains positional tetramer features from 250 sequences (15,000 sequences total). B. Structure plots of long (n=2,000) and short (n=2,000) Drosophila introns. Every sample is a single sequence. The topics are the same as the topics in Fig S5A. C. Top 10 enriched positional tetramers from topics 1 and 2 from the analysis of 5’SS of Drosophila introns. D. Sequence logo of the 5’SS from long Drosophila introns. E. Sequence logo of the 5’SS from short Drosophila introns. F. Sequence logo of positions 34 to 46 from long Drosophila introns. G. Sequence logo of positions 34 to 46 from short Drosophila introns. H. Distribution of topic 1 memberships of single sequences in Fig S5B. I. Distribution of the likelihood of generating sequences in Fig S5B by topics.

**Figure S6.**
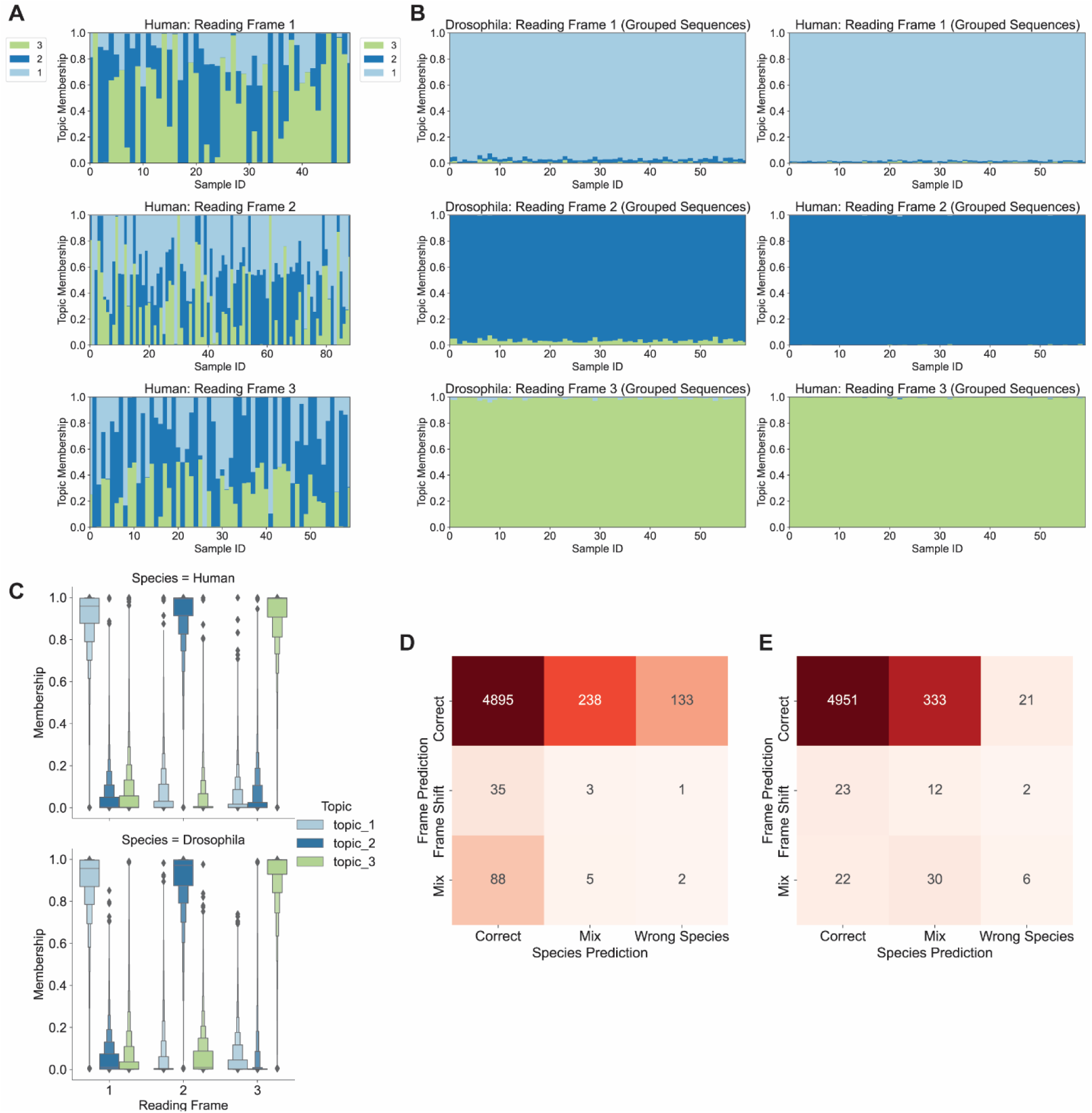
Misclassification from LDA models and generalization of human data fitted LDA model in predicting Drosophila reading frames. Reading frame classification was carried out based on topic distributions. Also, the ability to distinguish reading frames by human data fitted LDA model was tested on the Drosophila sequences. A. Topic memberships of human CDS sequences that are misclassified into incorrect reading frame tags. B. Structure plots of transforming human and Drosophila CDS sequences with human CDS-fitted LDA model. C. The topic distribution of sequences in Fig S6B. D. The confusion matrix of predicting reading frames and species for human CDS using the model in Fig 5A. E. The confusion matrix of predicting reading frames and species for Drosophila CDS using the model in Fig 5A.

**Figure S7.**
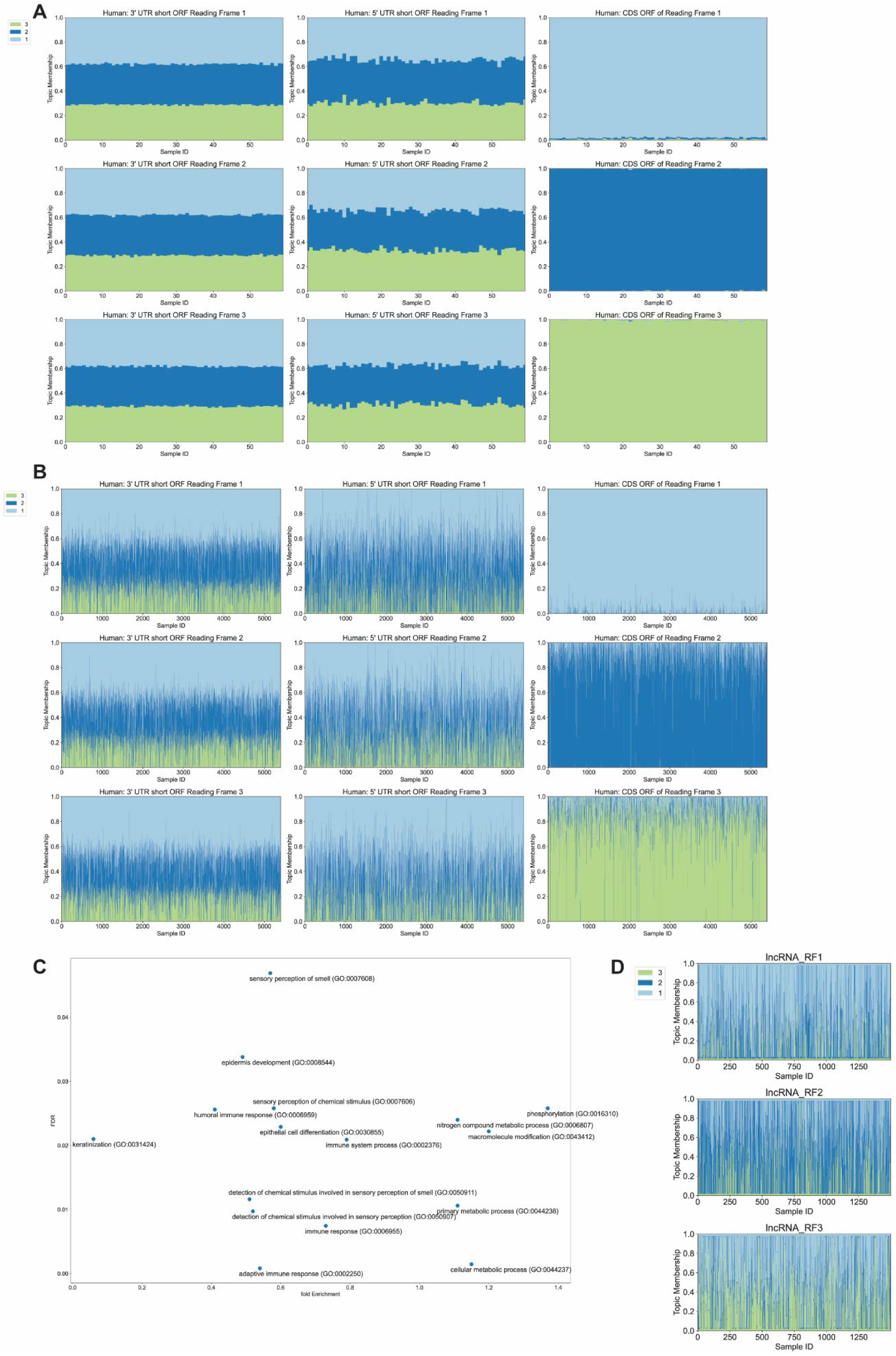
Analyzing short ORF in human UTR5 and lncRNA by CDS LDA model. Human data fitted LDA model was used to analyze short ORF from non-coding sequences. A. Structure plots of human small reading frames in UTR3, UTR5, and reading frames CDS. Data were transformed by the model Fig 4A. Each sample has hexamer compositions of 300 sequences (18,000 sequences total). B. Structure plots of human small reading frames in UTR3, UTR5, and reading frames CDS. Each sample is a single sequence from Fig S7A. E. Gene Ontology enrichment analysis of the smORF. F. Structure plots of human lncRNA small reading frames. Data were transformed by the model in Fig 4A. Each sample is the hexamer composition of a single sequence (1,500 sequences are shown).

